# Graded striatal learning factors enable switches between goal-directed and habitual modes, by reassigning behavior control to the fastest-computed representation that predicts reward

**DOI:** 10.1101/619445

**Authors:** Sean Patrick, Daniel Bullock

## Abstract

Different compartments of striatum mediate distinctive behavior-control modes, notably goal-directed versus habitual behavior. Normally, animals move back and forth between these modes as they adapt to changing contingencies of reward. However, this ability is compromised when dopaminergic drugs are used as reinforcers. These facts suggest that a set of biological variables, which make striatal decision making both highly plastic and uniquely sensitive to dopamine, contribute both to normal switches among modes and to the susceptibility for excessive habit formation when dopaminergic drugs serve as rewards. Indeed, data have revealed an impressive number of plasticity- and dopamine-related neural factors that vary systematically (with either increasing or decreasing gradients) across the rostral-ventral-medial to caudal-dorsal-lateral axis within striatum, the same axis implicated in switches among behavioral modes. Computer simulations reported here show how a dopamine-dependent parallel learning algorithm, if applied within modeled cortico-striatal circuits with parameters that reflect these striatal gradients, can explain normal mode switching, both into the habitual mode and returns to goal-directed mode, while also exhibiting a susceptibility to excessive habit formation when a dopaminergic drug serves as reward. With the same parameters, the model also directly illuminates: why interval and probabilistic reinforcement schedules are more habit forming than fixed-ratio schedules; why extinction learning is not (and should not be) a mirror image of acquisition learning; and why striatal decisions guided by reward-guided learning typically exhibit a highly sensitive tradeoff between speed and accuracy.

## Introduction

At any moment, typical invertebrate and vertebrate animals are capable of computing multiple representations of a given context, and of generating multiple behaviors within a context. Combined with the brute fact that trying to perform more than one representation-guided behavior at a time typically fails, unless each behavior recruits an effector system that is decoupled from the others, the omnipresence of multiple simultaneous representational and behavioral options implies that an animal’s waking life parses into a series of action-selection decisions. These decisions are of the form: Given my representations of context, which is most informative about the behavior I *should* generate *now*, and which of my alternative behaviors should that be? Because animals are subject to Darwinian evolution, the “should” question alludes to inclusive fitness, whereas the “now” question alludes to the non-stationarity (inconstancy across time) of contingencies relating decisions to proximal outcomes (e.g., safety, food, sex) that affect inclusive fitness. The best decision for a juvenile is often not best for an adult, the best foraging or mating decision in Spring is no longer the best in Autumn, etc. Thus even highly successful, oft-repeated, decisions may need to be avoided at later times, in service of inclusive fitness. Moreover, the optimal times/cues for switches of policy often cannot be established in advance. The timing of such switches can usually be improved by learning processes that reflect an interaction between an individual animal’s developing representational and behavioral abilities and its life experiences with non-stationary contingencies of rewards and punishments (Schultz, 2015).

Such learning and its role in behavioral switching have been studied across a broad spectrum of animals from arthropods to mammals. In recent years, solid empirical cases have been made for the thesis that the neural structures (Strausfeld & Hirth, 2013; Fiore et al, 2015) and neurotransmitters (Waddell, 2013) most vital for reward-guided decision making are at least highly analogous, if not homologous, across arthropods and mammals. In particular, both possess *forebrain nuclei* – in mammals, the *basal ganglia* – within which switches of decision-making are achieved by learning processes that are strongly influenced by dopamine (DA). Moreover, in arthropods and mammals, DA has now been implicated in learning guided by both rewarding and aversive events (reviews in: Waddell, 2013; Bullock, 2016). This implies that flexible switching that depends on reward-guided learning is truly ancient; this, in turn, has inspired debates about what the “oldest core” switching abilities are (e.g., Brembs, 2011; Perry et al., 2013), and how they may have been supplemented or enhanced in some lineages by evolution of further mechanisms.

Research on reward-guided decision-making in mammals supports a classification into two types of decision mechanisms: *model-free* versus *model-based* decision-making (Doya, 1999; Daw et al., 2005). In model-based decision-making, which is more cognitively advanced, the decision process depends, in some sense, on a mental anticipation, which in turn depends on a mental model/representation of at least one contingent relationship, namely the contingent relationship between an act and its probabilistic outcome. Model-free decision-making is not as cognitively advanced. By definition, a model-free decision process does not utilize any neural anticipation of an act’s outcome prior to selection of that act. As summarized in Figure 1, it is well established empirically (Yin & Knowlton, 2006; Yin et al., 2009) that protracted reinforced training often leads to a shift from a model-based to a model-free decision strategy.

**Figure 1.**
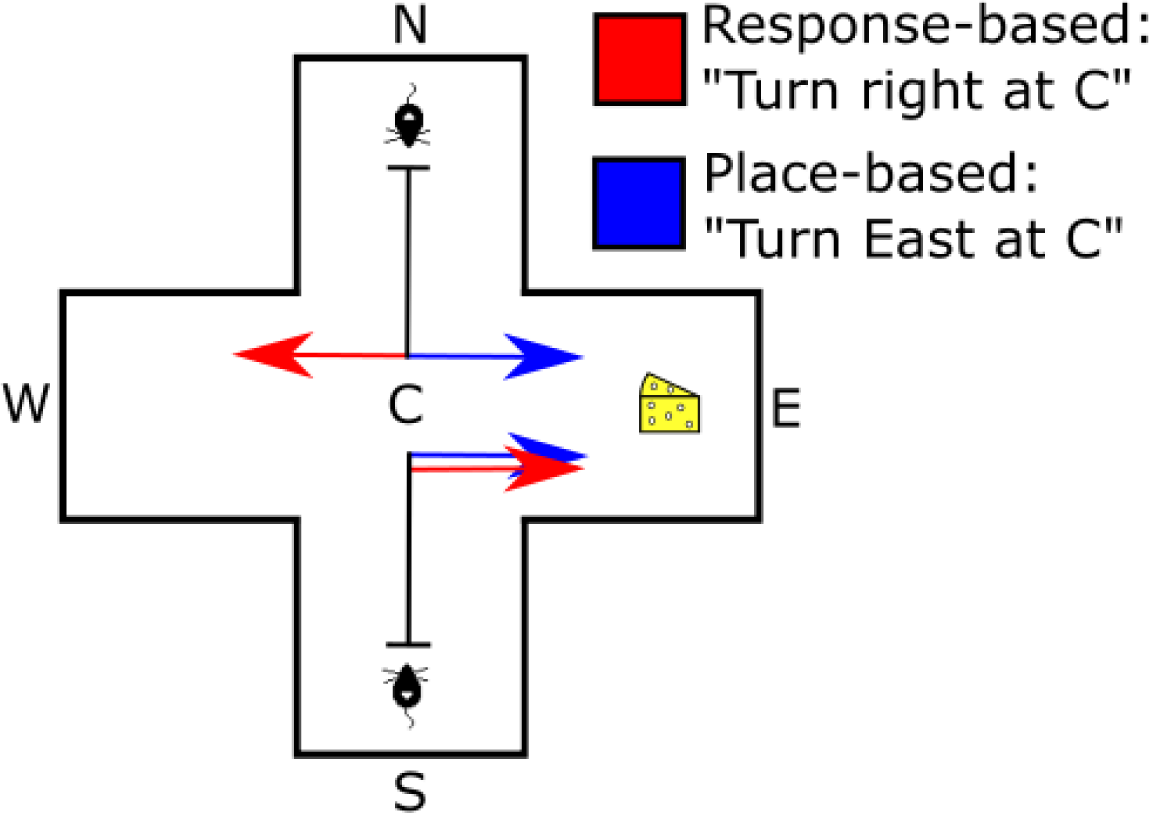
Protracted training leads animals to switch from a model-based (mental map or “place-based”) decision strategy to a model-free (“response-based”) strategy. On *training trials* in the classic plus maze, subjects start at the S (south) end and seek reward, which is always at location E (east). A turn decision occurs at central point C. On (rare) *probe trials*, subjects are instead placed at N to start. After modest training, probe trials reveal an ability to adjust decision making to correctly compensate for the novel starting point. This “place-based” strategy (“turn toward place E”) corresponds to the blue arrows. After much reinforced training, probe trials reveal habitual decision-making: in the example shown, the same subject now chooses a right turn regardless of the start position. The red arrows show that this response-based strategy results in an incorrect choice (no reward) when the subject is initially placed at point N.

Although the categorical distinction between model-free and model-based decision-making is well-grounded, and readily maps onto distinct neural circuitries, both varieties can be seen to occupy positions on a more fundamental continuum. Model-free decision-making is inherently faster than model-based decision making. This temporal aspect has been emphasized at least since the time when 1930’s behaviorists such as Hull – who emphasized model-free decisions mediated by reinforced or punished cue-action links – criticized cognitivists such as Tolman (who emphasized model-based decisions) for seeing animals as inherently slowed by cognitivists’ imputed requirement that animals access mental maps and compute outcome expectations before choosing to perform any learned actions. Furthermore, within the model-free part of the continuum, there is temporal differentiation because of the variable times needed to compute alternative representations from a given experiential stream. Even if an act is quickly selected based on contextual representations, without any pause to think ahead to the act’s consequences, alternate representations of a context take less or more time to compute. Similar temporal differentiation emerges within the model-based decision schemes, because computing the outcomes of acts can take less or more time, depending on the processes used.

Thus, a more fundamental distinguishing feature is the time taken to utilize whatever representations mediate a decision process. Broadly associated with this decision-time continuum is a continuum of decision quality. Many such associations between time and quality have been studied under the rubric of *speed-accuracy tradeoffs*. Animals without, or with, model-based decision-making, all face a tradeoff related to the time it takes to compute the representations on which decisions will be based. On average, more time allows better information integration, less noise-susceptibility, and higher quality decisions – whether for model-free or model-based decisions. Despite the average superiority of slower decisions, it is often observed that actors speed up their decision-making process over repeated encounters with a given choice scenario. This speed-up must reach a limit when the decision process is as fast as it can be, but evidence also suggests that it can stop well short of such limits, notably when decision quality verges on becoming insufficiently high to earn rewards with a high probability. That is, payoff statistics affect how an actor will tradeoff speed and accuracy.

The ubiquitous and ancient status of such tradeoffs warrant the inference that there are highly evolved biological mechanisms whereby behavior control is passed – by a reward-guided learning process – from slower, higher-quality, decision pathways to faster, lower-quality, decision pathways, but also, when contingencies so require, in the reverse direction. Important clues to the anatomical substrate for such mechanisms comes from research on the mammalian basal ganglia. The decision-speeding associated with a shift from model-based decision making to model-free decision-making involves a change in the sector of the striatum that mediates those decisions (Yin & Knowlton, 2006; Gruber & McDonald, 2012).

Whereas model-free decisions were mediated in the basal ganglia by dorso-lateral striatum (DLS), model-based decisions were mediated by more ventro-medial and dorso-medial parts of striatum (VMS/DMS). In this article, we report simulations of a neural circuit model that reveal learning and decision characteristics implicit in an interacting set of mechanisms that are suggested by data from anatomical, neurochemical, and neurophysiological studies of the basal ganglia and its major sources of input signals, notably cue representation-related signals from cerebral cortex, and reward-related signals from midbrain DA (dopamine) neurons. Preliminary simulations appeared in Patrick et al. (2014).

## Evidence for parallel learning processes in multiple brain regions that receive dopaminergic teaching signals

Research shows that learning is simultaneously occurring at many distinctive brain sites during experiential training trials (White & McDonald, 2002). Parallel learning is the rule, although in some cases seriality holds for short periods, due to the dependence of certain kinds of learning at site X on the emergence of representations at a site Y that projects to site X. An example of such transient seriality is the dependence of contextual fear learning on the hippocampal context representations that may take seconds to form after first exposure to a spatial setting (review in Krasne et al., 2011). There is evidence that parallelism holds for many kinds of DA-dependent learning, including those that occur in multiple distinct parts of the striatum, such as the (ventro-medial) nucleus accumbens, the (dorso-medial) caudate, the (dorso-lateral) putamen, as well as in striatum-like zones such as the central nucleus of the amygdala (CeA).

Some modelers (e.g., Keramati & Gutkin, 2013) have interpreted the late-in-training transfer of control from VMS/DMS to DLS during habit formation to imply seriality of learning epochs: they posit that substantial learning must occur in VMS/DMS before any learning begins in DLS. As possible support for their hypothesis, Keramati and Gutkin (2013) cite the discovery (Haber et al., 2000) that a spiraling linkage exists from VMS to DA neurons in VTA (ventral tegmental area) and SNc (substantia nigra, pars compacta), and from them to DLS, which in turn projects to SNc, but not back to VTA. However, contrary to the seriality hypothesis, this anatomy can be seen to ensure that learned changes in VMS are immediately reflected in both VTA and SNc, and thereby, in DLS. Moreover, a selective lesion to VMS leads to faster development of habitual (i.e. model-free, cf. Keramati & Gutkin 2013) control by DLS, whereas a selective lesion to DLS leads to indefinite prolongation of deliberative (i.e., model-based) control via VMS/DMS (Yin & Knowlton, 2006).

The faster development of DLS control in the absence of VMS/DMS and related phenomena (reviewed in Burton et al., 2015) are incompatible with the serial model of Keramati and Gutkin (2013). Given that reward-related dopamine bursts occur in both VMS/DMS and DLS virtually from the beginning of training (reviews in Schultz, 1998, 2015), and that the other pre-conditions for cortico-striatal synaptic plasticity identified *in vitro* or *in vivo* (review in Gurney et al., 2015) are also satisfied in both striatal zones, a more natural interpretation – adopted here – is that learning occurs in parallel, virtually from the start of training, but that, due to factors noted below, the VMS normally outcompetes the DLS for behavior control until many learning trials have accumulated.

In particular, our neural circuit model explains delayed transfer of control, despite parallel learning, by incorporating two abstract computational features supported by numerous neurobiological factors that vary in a graded way along the axis running from the ventro-medial to the dorso-lateral poles of the striatum. The emergent computational features we modeled are that, under normal physiological conditions, (1) learning rates are higher towards the ventro-medial than the dorso-lateral pole and (2) asymptotic learned synaptic weights are higher towards the dorso-lateral than the ventro-medial pole. These factors, with empirical references, are listed in Table 1, and Figure 2 explains how the first factor – less DAT (DA transporter) in VMS/DMS than DLS – contributes to faster initial learning in VMS/DMS. We next discuss how the factors listed in Table 1 may relate to these two features of parallel learning across striatal compartments.

**Table 1.**
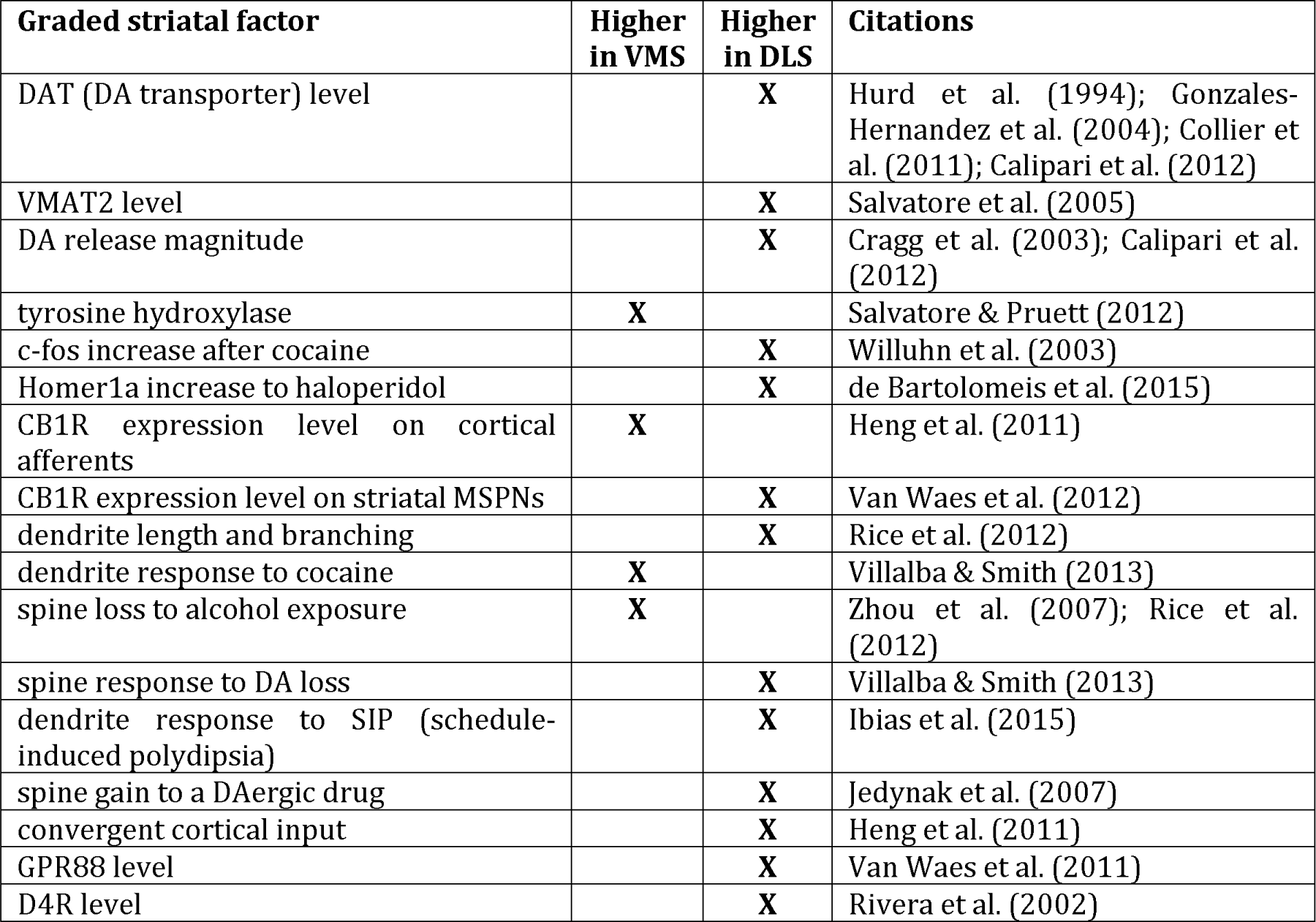
Gradients of learning-related factors along a behavior control axis of the striatum.

**Figure 2.**
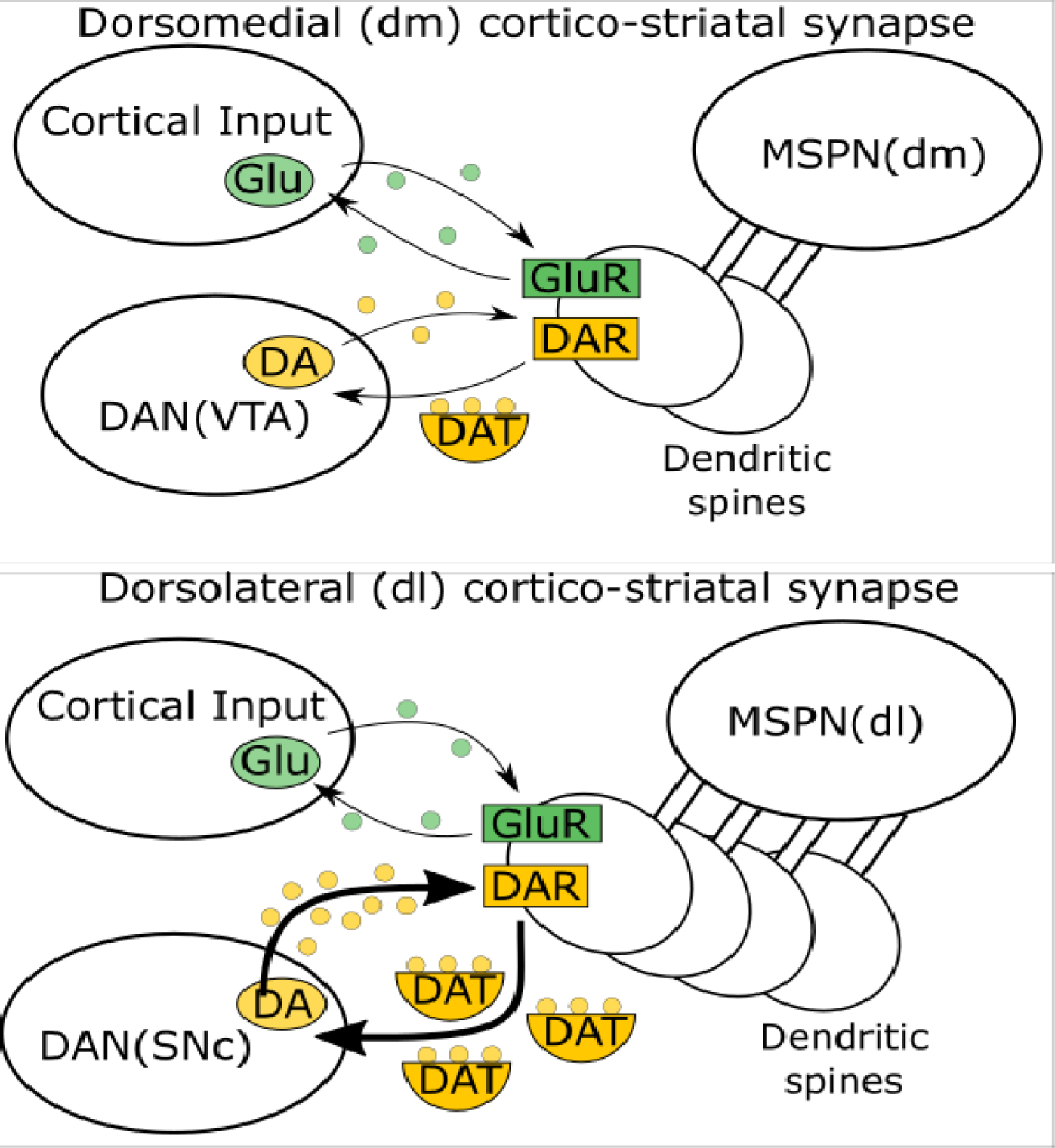
Effects of dopamine (DA) differ between the dorsomedial (dm) and dorsolateral (dl) zones of the striatum. Axons projected to striatum by DA-ergic neurons (DANs), whose somas are in either the substantia nigra pars compacta (SNc) or ventral tegmental area (VTA), release DA at dendritic loci near synapses formed between cortical pyramidal neuron axons and medium spiny projection neurons (MSPNs) of the striatum. Following release, DA is cleared from the synapse by DA transporter (DAT) and returned to the DAN’s axon terminal. DA modulates long-term potentiation and depression (LTP and LTD) of the effects that cortical inputs have via glutamate receptors (GluRs) on striatal MSPNs. DA modulates LTD or LTP via DA receptors (DARs) on MSPN dendritic spines. In the dorsolateral zone, DA release sites are denser, so reward-induced DA releases are denser (more yellow dots), but DAT is also denser. Thus DA reaches more synaptic growth sites quickly (after short diffusion-times), and faster clearance of DA by denser DAT implies a shorter time for DA to act at each locus to promote spine/synapse formation. The

## Evidence for faster learning in ventro-medial than dorso-lateral striatum

There is abundant evidence that concentrations of DAT, the dopamine (DA) transporter, increase along the ventro-medial to dorso-lateral axis within the striatum (Gonzalez-Hernandez et al., 2004; Wickens et al., 2007; Collier et al., 2011; Calipari et al., 2012). Because the rate at which DAT removes previously released DA is concentration-dependent, this implies that burst-released DA remains available to promote learning for briefer intervals in DLS (dorso-lateral striatum) than in VMS (ventro-medial striatum), and thus that learning rate per trial is slower at points nearer the DLS pole than at points nearer the VMS pole. The faster learning in VMS/DMS than DLS, by itself, creates an advantage for initial learned behavior control via VMS/DMS rather than via DLS. This is consistent with the behavioral hypothesis of Dickenson et al. (1995) that learning rate is higher for act-outcome learning – now known to be mediated by DMS – than for stimulus-response learning, now known to be mediated by DLS. Further, there is strong evidence that the DAT gradient is predictive. If DAT is higher in DLS, and it matters, then DLS should be more strongly affected by DAT blockade than VMS/DMS. Indeed, cocaine, a DAT blocker, produces a much larger increment in c-fos expression in DLS than in other zones of striatum (Willuhn et al., 2003).

## Evidence for a higher learned synaptic weight asymptote in dorso-lateral than ventro-medial striatum

The model’s second circuit feature implies that feature one – the learning rate advantage held by VMS/DMS – can eventually be overcome by greater accumulated learning in DLS. That this might be true is suggested by the finding that high levels of practice often cause a shift from VMS or DMS to DLS control of behavior (Yin & Knowlton, 2006). Evidence also suggests that the asymptotic strength of learning-adjusted cortico-striatal connections is higher at the DLS pole than at the VMS pole under normal physiological conditions. For example, more spines create more sites for synaptic contacts, and higher learned synaptic weights (Araya, 2014; Araya et al., 2014; Mancuso et al., 2014; Forsyth & Lewis, 2017). Some have reported that striatal MSPNs’ (medium spiny projection neurons’) spine densities (cf. Figure 2) increase from shell to core along the ventromedial to dorsolateral axis within ventral striatum (Meredith et al., 1992; 2008).

Although some have failed to find such a gradient (Rice et al., 2012), spine density, per se, is a very labile variable that changes with learning history, because events that induce learning do so in part by inducing new spine formation. A more experience-invariant statistic is dendritic length and the number of branches, which are both higher for DLS MSPNs than for VMS MSPNs (Rice et al., 2012). This creates a larger surface area from which spine growth can be induced during learning involving DLS MSPNs. Moreover, research has shown that dendrite growth and spine formation depend jointly on DA-ergic and cortical inputs to striatum (Penrod et al., 2015), and DLS (relative to VMS) has higher levels of convergence from multiple cortical areas (Heng et al., 2011), a higher density of DA-ergic fiber terminals, and a greater amount of DA released per volume of tissue (Calipari et al., 2012). These trends are consistent with the finding that reduced DAergic innervation of DLS, following loss of SNc DA neurons in Parkinson’s Disease, results in markedly reduced DLS spine densities, and reduced learning ability (Villalba & Smith, 2013). Consistently, a drug that increases DA release, methamphetamine, induces increases of spine density in DLS, but decreases of spine density in DMS, changes that are associated with faster shifts from model-based to model-free (habitual) behavior (Jedynak et al., 2007). The higher basal level of dendritic length/branching, combined with greater convergence of cortical afferents, create a greater potential for DA-influenced spine formation in DLS. This greater potential, and the normal correlation of spine density with synaptic efficacy, imply that, *if DA-gated learning is protracted enough*, then efficacies of cortico-striatal synapses in DLS can eventually exceed the maximum achievable efficacies in more ventro-medial zones of striatum.

The ‘if’ condition is emphasized because of the well-established principle (cf. Schultz, 2015) that the burst DA signals that gate learning in striatum typically reflect RPE (reward prediction error), which is largest at the onset of learning, but then progressively shrinks if and as the system learns to use context-related representations to predict the timing and magnitude of reward. Given that principle, it is important to also note that a significant burst DA signal can persist *indefinitely* in typical real-world cases, because reward typically is not fully predictable. In particular, reward is not fully predictable whenever action-outcome contingencies are probabilistic, and whenever the timing of reward-delivery following an instrumental action is variable. This observation dovetails with findings (Yin & Knowlton, 2006) that transitions to habitual control via DLS are more likely to occur on interval and variable-ratio reinforcement schedules – for which it’s impossible for an animal to predict that a current response will be rewarded – than on fixed-ratio schedules, for which exact or approximate prediction is possible. Furthermore, there is direct evidence that when CRfs (conditioned reinforcers) are response-contingent, then CRf-related burst DA events in DLS, which receives DA released by fibers from the SNc, persist at a much later point in the learning process (i.e., after a greater number of CRf-contingent training trials) than CRf-related burst DA events in VMS, in which DA is released by fibers from the VTA but not the SNc (Ito et al. 2000; 2002). Using cocaine as reward, Willuhn et al. (2012) showed that during training, the response-contingent DA release in DLS increased as the release in VMS (i.e. NAcc core) decreased. The reliably later waning in DLS than VMS can be plausibly attributed, at least in part, to slower learning (due to higher DAT, as noted above) in the cortico-DLS-SNc pathways that are responsible for learned suppression of reward-related burst firing in SNc (and DA release in DLS) than in the cortico-VMS-VTA pathways that are responsible for learned suppression of such DA bursts in VTA. Here the “are responsible” attributions are warranted by simulations of learned DA neuron RPE responses within local-circuit models (e.g., Tan & Bullock, 2008; Chorley & Seth, 2011; Schultz, 2015) of the known loops that link striatum and dopamine cells.

What explains the waxing of the DA release in DLS during the training epoch, as examined by Willuhn et al. (2012)? This effect may be explicable in terms of the “Haber spiral” (Haber et al., 2000), which relates the VMS vs. DLS distinction to different DA neuron pools (in VTA vs. SNc) that differ in their inputs and outputs. For example, whereas VMS neurons that make direct inhibitory synapses on VTA DA neurons may be responsible for waning of the VTA DA signal delivered to VMS, other VMS neurons may, via direct synapsing on medial SNc GABAergic interneurons (INs), actually disinhibit medial SNc DA neurons projecting to DLS (see Haber et al., 2000, Figure 12; Watabe-Uchida et al., 2012). The latter cells’ activity would wax larger as learning in VMS proceeded.

Indeed, there is already some evidence that such distinctions among DA neuron pools help explain protracted occurrences of DA bursts in DLS after such bursts have waned in VMS. So far, we have emphasized only two axes of the mammalian striatum: the dorsal-ventral and the medial-lateral. Although these receive the most research attention, also pertinent is the rostral-caudal, or anterior-posterior axis (which is partly confounded, because the most ventral-medial parts of striatum are present in rostral sections, but disappear from caudal sections). Several studies in humans have associated model-free/habitual control with the posterior putamen (Tricomi et al., 2009; de Wit et al., 2012). This suggests that it may be more accurate to speak of a rostral-ventral-medial to caudal-dorsal-lateral axis (see also Voorn et al., 2004). Moreover, a recent study in mice (Menegas et al., 2015) discovered that the DA neurons in lateral SNc that project to “tail” or caudal striatum are unique – relative to DA neurons that instead project to more rostral-ventral parts of striatum – in two ways: they receive proportionately fewer monosynaptic inhibitory inputs from ventral striatum, and they receive proportionately more numerous monosynaptic inputs from paraSTN/STN (subthalamic nucleus), the GPe (external globus pallidus) and the ZI (zona incerta). As Menegas et al. (2015) also noted, the relative lack of inhibitory afferents from ventral striatum warrants the expectation that this unique group of DA neurons will not show the same RPE-related behavior (i.e., a waning response to predictable reward) as other DA neurons that project to more rostral-medial-ventral parts of striatum.

Such observations are consistent with the parallel learning hypothesis and the thesis that typical DA cells responsible for DLS habit learning continue to burst long after such bursts have waned in DA cells that control learning in other parts of striatum. Further, the greater inputs from GPe, paraSTN/STN, and ZI are likely to be very pertinent to understanding the protracted skill learning that occurs after initial response acquisition. Lingawi & Balleine (2012) reported on a paradigm in which the transition to habitual control depended on an intact CeA (itself striatum-like; see Swanson, 2000 and Swanson, 2006), and Chometton et al. (2017) recently reported strong reciprocal links between CeA and paraSTN in rodents. Furthermore, progressive skill learning often involves not only response-speeding but also changing the manner of performance, by amplifying helpful response components and pruning away response components that reduce an action’s success (Buitrago et al., 2004; Yin et al., 2009). In this phase, the cerebellum is also strongly engaged, and is linked to all three areas noted in the report by Menegas et al.: there is a projection from primate ACC area 24 (Schmahmann & Pandya, 1997), implicated in performance-error monitoring, via ZI to the olivary nuclei (Saint-Cyr & Courville, 1981), which are the sole source of performance-error signals relayed via climbing fibers to the cerebellum; there is a projection from sensori-motor STN to pontine nuclear sources of mossy fibers to the cerebellum (Bostan et al., 2010); and there is a projection from the cerebellar dentate nucleus via an intralaminar (central-median) nucleus of the thalamus, and the dorso-lateral striatum, to GPe (Ichinohe et al. 2000; Hoshi et al. 2005).

Another difference between VMS and DLS (see Table 1) is a reversed pattern of expression of CB1Rs (cannabinoid type 1 receptors): in VMS, CB1Rs are denser on cortical afferents (Heng et al., 2011) whereas in DLS, CB1Rs are denser on MSPNs (Van Waes et al., 2012). Wang et al. (2006) showed that CB1Rs regulate DA- and ACh-dependent plasticity of glutamatergic cortico-striatal synapses, and Wu et al. (2015) showed that activation of D1Rs in direct (“GO”) pathway MSPNs could mask CB1-dependent LTD, whereas activation of D2Rs could boost CB-LTD in indirect (“NOGO”) pathway MSPNs. Consistently, Martin-Garcia et al. (2016) found that the associative aspect of learned cocaine self-administration depended more on CB1Rs on glutamatergic fibers from cortex than on CB1Rs expressed in striatal MSPNs. The greater expression of Homer1a in DLS than in VMS (de Bartolomeis et al. 2015) following haloperidol (Table 1), provides further evidence of D2R-related plasticity differences between VMS and DLS, because haloperidol selectively blocks D2Rs, and Homer1a is associated with synaptic plasticity mediated by restructuring of postsynaptic densities.

The final two rows of Table 1 list two other receptor types with a gradient distribution across striatum: D4R and GPR88. The D2R-family D4 receptors appear in spines and dendrites of MSPNs, with greater expression in striosomes than matrix, and greater expression in DLS than VMS (Rivera et al., 2002) in most species examined (rodents and primates, but not cats). D4Rs are involved in control of mu-opioid receptor expression (Gago et al., 2007), and certain genetic variants of the D4R have been implicated in gambling habit formation (Eisenegger et al., 2009) in PD patients. The D4R presence in striosome MSPNs that project to midbrain DA neurons is suggestive, because in general habit formation has been associated with abnormally elevated DA release, diminished processing of response costs by DA neurons, and reduced negative RPEs. The G-protein-coupled receptor GPR88 has a higher expression in DLS than VMS (Van Waes et al., 2011), and its lack leads to deficient spine growth on D2R expressing MSPNs. Meirsman et al. (2014) found that mutants lacking GPR88 showed non-habituating hyperactivity, enhanced stereotypy, and a deficit in rotorod habit learning, which depends on recruitment of D2R-expressing MSPNs in DLS across multiple training sessions (Yin et al., 2009). Consistent with that DLS deficit, the GPR88-lacking mutants also showed enhanced dependence on spatial goal-directed (model-based) behavior, mediated by hippocampus and VMS/DMS.

In summary, dendritic and receptor gradients spanning from VMS through DMS to DLS imply learning differences at various points between the high-versus low-density poles of those gradients. The weight of the evidence supports the hypothesis that, under normal physiological conditions, asymptotic learned cortico-striatal weights in DLS are greater than in VMS/DMS. This hypothesis, a key assumption of our model, is contrary to the model of Keramati and Gutkin (2013), which instead supposed that asymptotic weights in DLS become greater than those in VMS/DMS only under the influence of addictive drugs. In the literature (review in Gruber & McDonald, 2012) VMS is associated with learned control of unlearned orientation, approach, and withdrawal behaviors, whereas DMS and DLS are associated with learned contextual control of learned instrumental behaviors, such as forelimb reaching-grasping. To focus on how stimulus-cued instrumental behaviors can be brought under learned control via two alternate pathways, we will now address transitions of control from DMS to DLS, and the reverse.

## How parallel learning and two modeled asymmetries explain delayed transitions to habitual control

The effects, on parallel reward-gated synaptic changes, of two modeled circuit factors – learning rate and asymptotic synaptic weight sizes – are schematized in Figure 3. If the relative magnitude of the synaptic weights in DMS versus DLS is the *principle* determinant of which region mediates the decision on a given trial within a protracted series of decisions that are made under a stationary contingency linking action to reward, then Figure 3 in effect illustrates how the two factors imply a switch between early control by DMS and later control by DLS. However, other factors, including the time needed to compute a cortical representation, and so access its cortico-striatal weights, can also affect such competitive decisions. In our computational model as specified below, we assume, consistent with the main trend in mammalian data, that cortical representations that project to DLS, described here as ‘primary’ are computed faster than the cortical representations, described here as ‘secondary’, that project to DMS. These are meant to be relative terms. The effect is that the learning process assigns control earliest to slower-computed, presumably more accurate and integrative, secondary representations, but may later pass control to faster-computed, primary representations.

**Figure 3.**
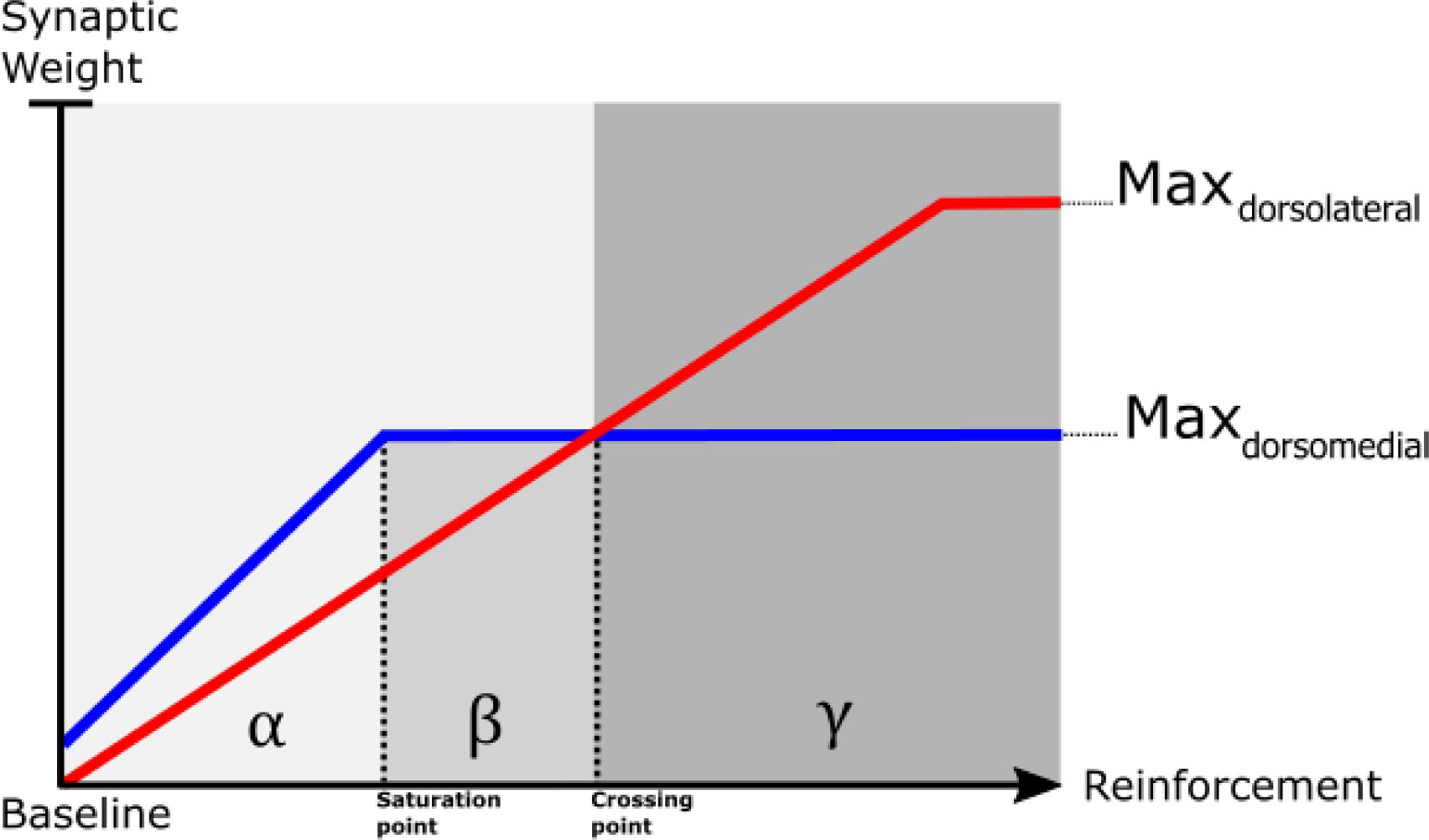
Two factors that affect control switches between dorsomedial (DMS) and dorsolateral (DLS) striatum. Pictured on the vertical axis is the synaptic weight, a normalized measure of cortico-striatal synaptic efficacy that depends jointly on the excitatory effect of the direct (“GO”) pathway and the inhibitory effect of the indirect (“NOGO”) pathway, by which striatum controls behavior. The value of this weight is adjusted by DA-dependent “reinforcement” signals induced by receipts (or omissions) of rewards. Accumulated effects of net-positive reinforcement signals (shown on the horizontal axis) may increment the weight to its maximum (Max) value. Solid lines correspond to the best-case weight trajectory hypothesized to be possible for either DMS (blue line) or DLS (red line). Note that the DMS weight (blue line) has a higher slope but lower Max value than the DLS weight (red line). These hypothesized differences are based on regional differences in DA release, DAT, and both resting and potential spine density. Three decision-zones emerge (gray background). In the leftmost zone, marked ‘α’, a higher synaptic weight in the DMS results in faster accumulation (and threshold crossing) and therefore behavioral control by the “model-based control” region (DMS). In contrast, in the γ zone, entered only after sufficient reinforcements have pushed the DLS weight past a crossing point (the Max value of the DMS weight, Max_dorsomedial_), control is preempted by the “model-free” region (DLS). The intermediate decision zone, denoted ß, lies between the red-blue crossing point and the point at which the DMS synaptic weights saturate (i.e., no longer respond to positive reinforcements, due to limitations on spine density). Judging by only the relative weight values, the ß zone would entail exclusive control by DMS. However, cognitive processing creates greater time-delays (not depicted) for cortical inputs to DMS than to DLS, and thus may delay the onset of evidence accumulation in DMS by enough to allow DLS to sometimes gain control in the ß zone.

## Distinct pathways and learning rates for learned promotion vs. learned demotion of behavioral options

Because contingencies linking actions to rewarding outcomes are highly non-stationary in nature, a key question is how well the factors schematized in Figure 3 work within a circuit that enables adaptive control switches between alternative actions whenever changes of contingencies create new rank orderings of the expected values of those alternative actions. To adapt to such changes, a new rank ordering of synaptic weights must be created by some constellation of reward-guided learning processes. One such process is that of incrementing and decrementing synapses that *promote* action, both already implicit in Figure 3. However, there are many reasons to also include a parallel, synergistic, process that involves decrementing and incrementing synapses that *oppose* the same action. One reason is that, in vertebrates studied to date (Reiner, 2009), the basal ganglia are characterized by two opposing pathways for behavior control, a “direct pathway” that promotes action, and an “indirect pathway” that opposes action. In the striatum, these pathways utilize opposing sub-classes of the striatum’s principal neuron type, the medium spiny projection neuron (MSPN). MSPNs that express type 1 DA receptors and co-release GABA and SP (substance P), the D1-MSPNs, are so embedded in the circuit that their activation promotes an action’s selection; whereas activation of nearby MSPNs that express type 2 DA receptors and co-release GABA with ENK (enkephalin), the D2-MSPNs, opposes the same action’s selection.

In mammals, it has also been shown (review in Bullock, 2016) that burst releases of DA have opposite effects on the incrementing or decrementing of excitatory synapses onto these two classes of MSPN. This allows such synaptic adjustments to be synergistic. Two notable benefits of this synergistic solution are: it makes the learning process more robust to any mutations that degrade one or the other, but not both; and it implies that the process of behavioral extinction (learned demotion of a formerly preferred action) is not the mirror image of initial behavioral acquisition. Notably, whereas initial acquisition may often solely involve incrementing of synaptic weights that promote an action, extinction will normally involve a mixture of two processes: the decrementing of synapses that promote an action and the incrementing of synapses that demote that action. One consequence is that an action can be demoted more quickly if it no longer leads to reward, a contingency reversal that is often context-, or time/season-, dependent (Bouton et al., 2012; Trask et al., 2017). A related consequence is that much of the synaptic change that enabled acquisition need not be reversed during extinction, which, again, may induce context-specific learning. Instead, the action’s expression will be temporarily blocked by the net effect of incremented synapses that demote the action. Besides being critical to achieve basic circuit realism when modeling basal ganglia (as well as other parts of the brain, e.g., cerebellum), the concept of *synergistic adaptive opponent pathways* has significant explanatory power with respect to learned behavior control.

Notably, behavioral data usually show a faster-reacquisition phenomenon sometimes called “savings”: *re-acquisition*, upon reinstatement of an action-reward contingency (following a period of non-reward, e.g., in an uncued extinction protocol), is typically much faster (requires many fewer training trials) than *initial acquisition*. That these consequences would be highly adaptive phenotypes in all animals helps explain the ancient status of the direct and indirect pathways in vertebrate basal ganglia (Reiner, 2009). In the light of such neurobiological and behavioral data, the simulated model includes two kinds of striatal neurons subject to opposite rules for reward-guided changes to the weights of their excitatory synaptic inputs: D1-MSPNs and D2-MSPNs. These are referred to in the context of the neural circuit model, depicted in Figure 4, as “GO” and “NOGO” neurons, based on their respective tendencies to either facilitate or inhibit behavior. *Within* a given modeled region (DMS vs. DLS), we assume that the GO and NOGO weights have the same asymptote. However, consistent with some prior synaptic learning models (e.g., John et al., 2013), which simulated the asymmetry between acquisition vs. extinction, and findings of fast removal of learned inhibition during reacquisition learning, we assume that learning rates of cortico-striatal synapses are greater (by a factor of 5) for NOGO MSPNs than for GO MSPNs of the same region.

**Figure 4.**
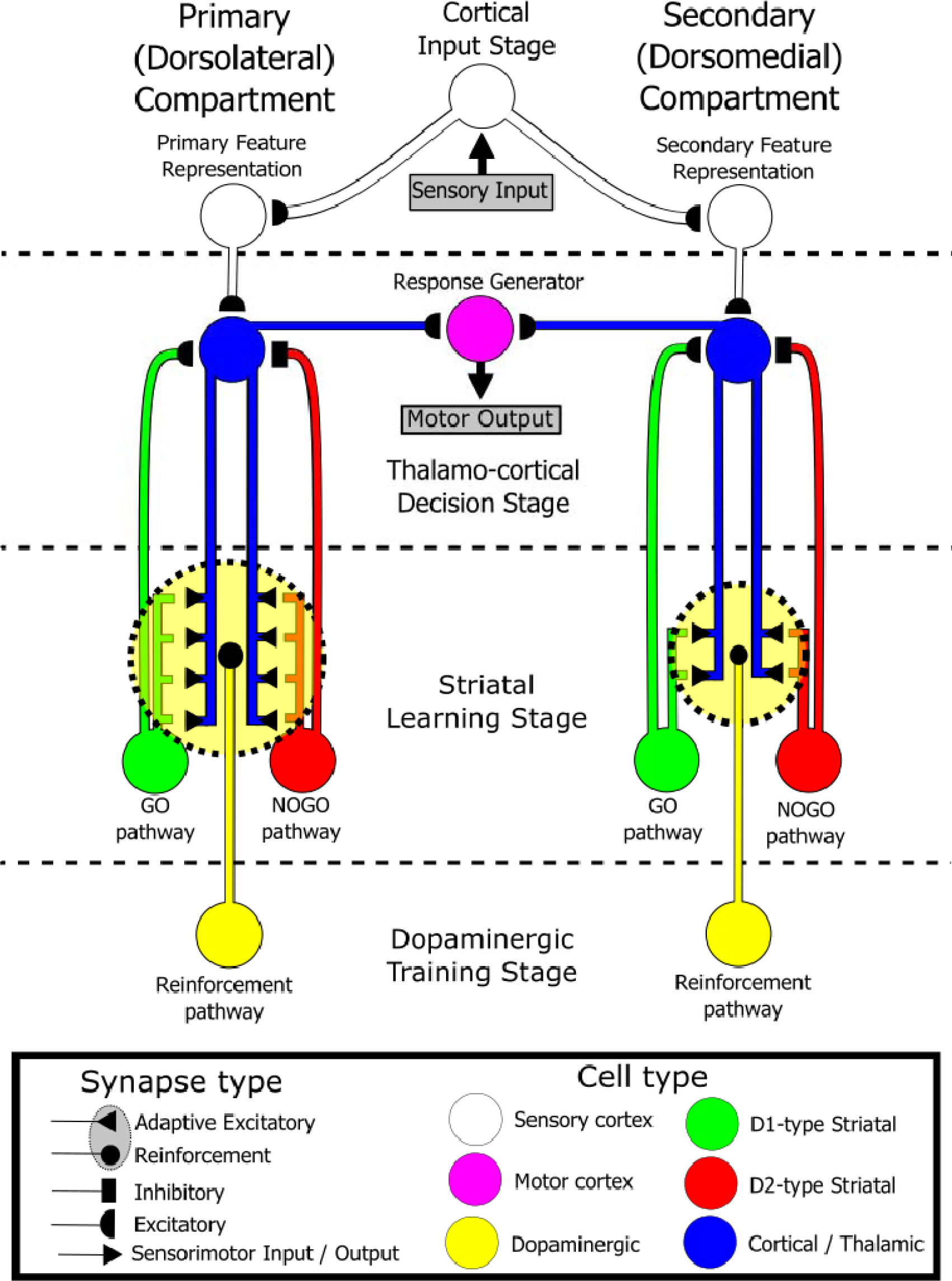
Neuronal constituents of a two compartment model of how reinforcements guide switches of control among alternative representations. Model elements constituting either a dorsolateral (left) or a dorsomedial (right) striatal subregion (and associated cortico-thalamic sectors) are depicted side-by-side as parallel model “compartments”. The dorsolateral subregion corresponds to the “primary” compartment, so-called because it receives signals from cortical neurons that represent quickly-computed, relatively “primary” features of stimuli. These give model-free representations of the context for decision, whereas representations available to the secondary/dorsomedial compartment take longer to compute and may be model-based representations. Within each of these compartments are multiple parallel instances of the cortico-basal ganglia loop pictured above, with each loop corresponding to a distinct learnable contingency relating a stimulus-feature to a response-generator (e.g., “Stimulus 1 -> Response 1”). In accord with the differences between DMS and DLS noted in Figures 2 and 3, and Table 1, the potential for higher asymptotic synaptic weights in primary than secondary compartments is schematized by additional synaptic contacts (black terminals of blue cortico-striatal fibers) in the primary compartment. Yellow ovals (bounded by thick, dashed lines) indicate the modulatory effects of diffusing dopamine on plasticity of cortico-striatal projections that reach both GO MSPNs (green) and NOGO MSPNs (red).

The aim of the simulations that follow is to illustrate how the model explains autonomous learned shifts among control modes, while relying on the four principles introduced above: (1) parallel learning in DMS and DLS compartments; asymmetries across striatal compartments in dopamine-dependent (2) learning rates and (3) synaptic weight asymptotes; and (4) faster NOGO than GO learning. Furthermore, monotonic waning of reward prediction errors (RPEs) across learning trials is incorporated to emulate the main trend of observed changes of dopamine release, with faster waning in the model’s DMS than its DLS compartment. To model the observed initial bias toward DMS over DLS control in biological learning, cortical-striatal synaptic weights onto D1-MSPNs are initialized with slightly larger values in the DMS compartment. Because the model is implemented in a way that remains agnostic to cortical processing differences other than the fundamental difference that some representations take longer to compute than others, none of the results reported here depend on assumptions about specialized features of representations computed outside striatum, e.g., in neocortex, hippocampus, or basolateral amygdala.

All simulations were made stochastic by adding noise at the cortical decision stage. The simulations illustrate how the computational model (specified in Appendix A) can be used to implement a wide range of behavioral learning protocols, and also demonstrate the emergence within the model of a number of key behavioral phenomena including: autonomous reward-guided transfers of control; savings of acquisition-learning across extinction episodes; faster relearning across repeated contingency reversals; and the typical speed-accuracy tradeoff between alternative decision strategies. Parametric simulations establish that these behaviors are robust and attributable to the model’s operation in parameter ranges that ensure the four listed principles.

All of these simulations examine the consequences for decisions of the effects of burst dopamine-release events on learning. With one exception, they do not simulate any direct effects of dopamine on the rapid time scale of decision processing. The sole exception uses reversal learning to illustrate how the so-called “uncertainty response” of dopamine cells (Fiorillo et al., 2003), is above-baseline DA-release distinct from burst DA-release, will have a much larger effect in low-DAT regions of striatum than in the high-DAT DLS. Thereby, the DAergic uncertainty response, which may be ancient because it can be computed subcortically in VTA/SNc (Tan & Bullock, 2008a, 2008b; Schultz, 2015) using MSPN co-release of GABA and SP (which is ancient in vertebrates; review in Reiner, 2009), can promote use of secondary representations, and model-based control, during conditions of uncertainty, consistent with proposed abstract rules for rational switching (Daw et al., 2005). Finally, although the model was formulated to explain learning guided by arbitrary rewards, its mechanisms directly imply enhanced habit formation when DAergic drugs serve as rewards, as illustrated with one simulation.

## Results

Using the model to simulate overtraining and reversal learning protocols demonstrates the efficacy of control transfer. Learning-dependent transfers of control were repeatable, and preserved prior learning. Figure 5 allows comparison of the model’s initial acquisition of a conditioned response with its reacquisition of that conditioned response after a prolonged extinction block. In both the initial acquisition and reacquisition blocks, the model acquires the conditioned response to within 90% accuracy. Reacquisition, during resumption of the contingency after extinction, occurs much faster than the initial acquisition. This reproduces the longstanding reinforcement learning phenomenon of rapid reacquisition (Figure 5c), also known as “savings”, which appears following extinction in both Pavlovian and instrumental learning paradigms (Kehoe & Macrae, 1997; Bouton et al., 2012; John et al., 2013; Crossley et al., 2016).

**Figure 5a.**
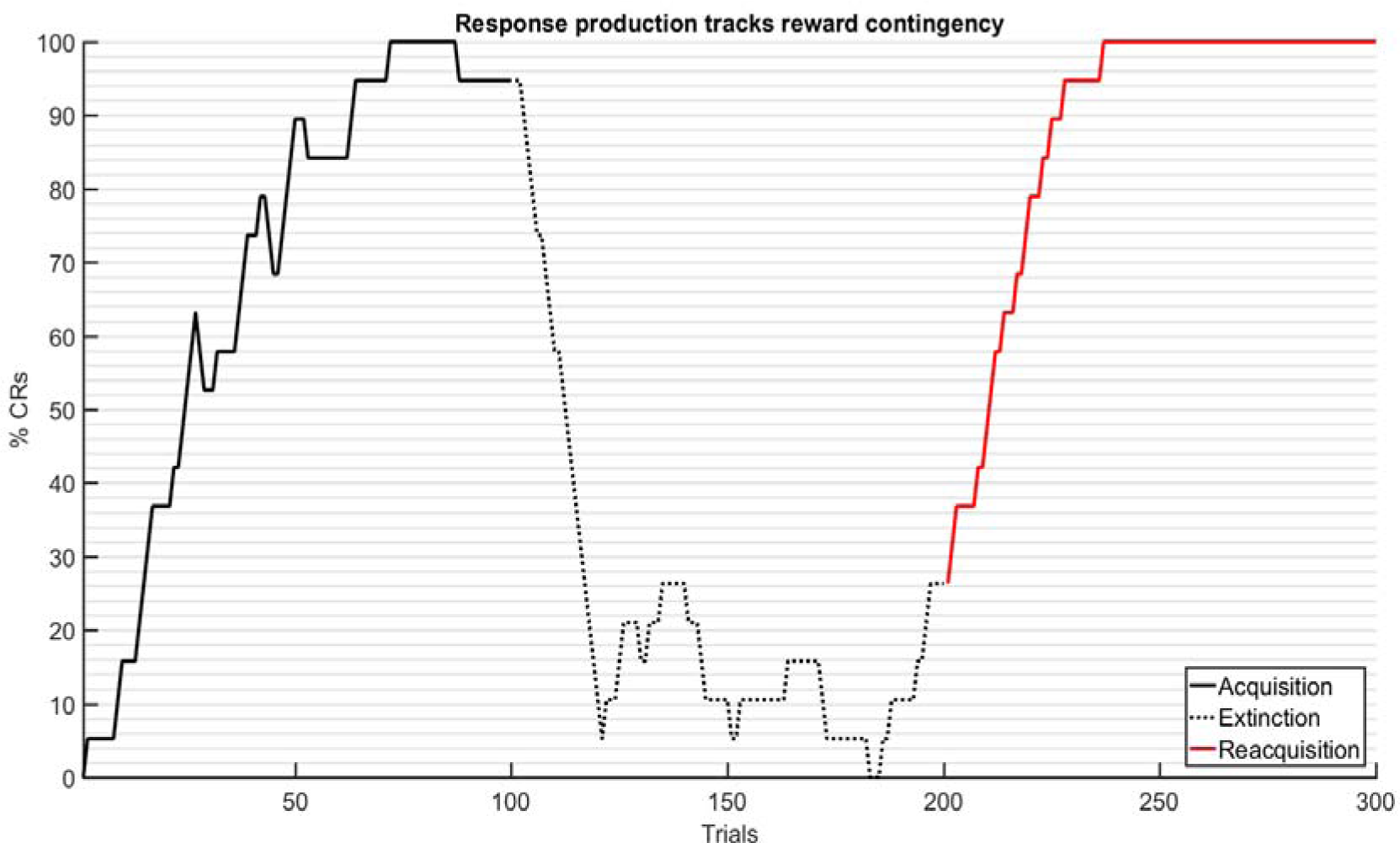
Acquisition, extinction and reacquisition. Twenty copies of the model (each with independent noise added at the cortical decision stage) were rewarded for choosing response 1 during trials 1-100 (acquisition, solid black line) and 201-300 (reacquisition, red dotted line), but were subjected to an extinction phase, in which no response was rewarded, from trials 101-200 (black dotted line). For every simulation and the average of all simulations (plotted here) the percentage of chosen responses that were correct (and thus rewarded) rose above 90% in the acquisition and reacquisition phases, but fell below 10% late in the extinction phase.

**Figure 5b.**
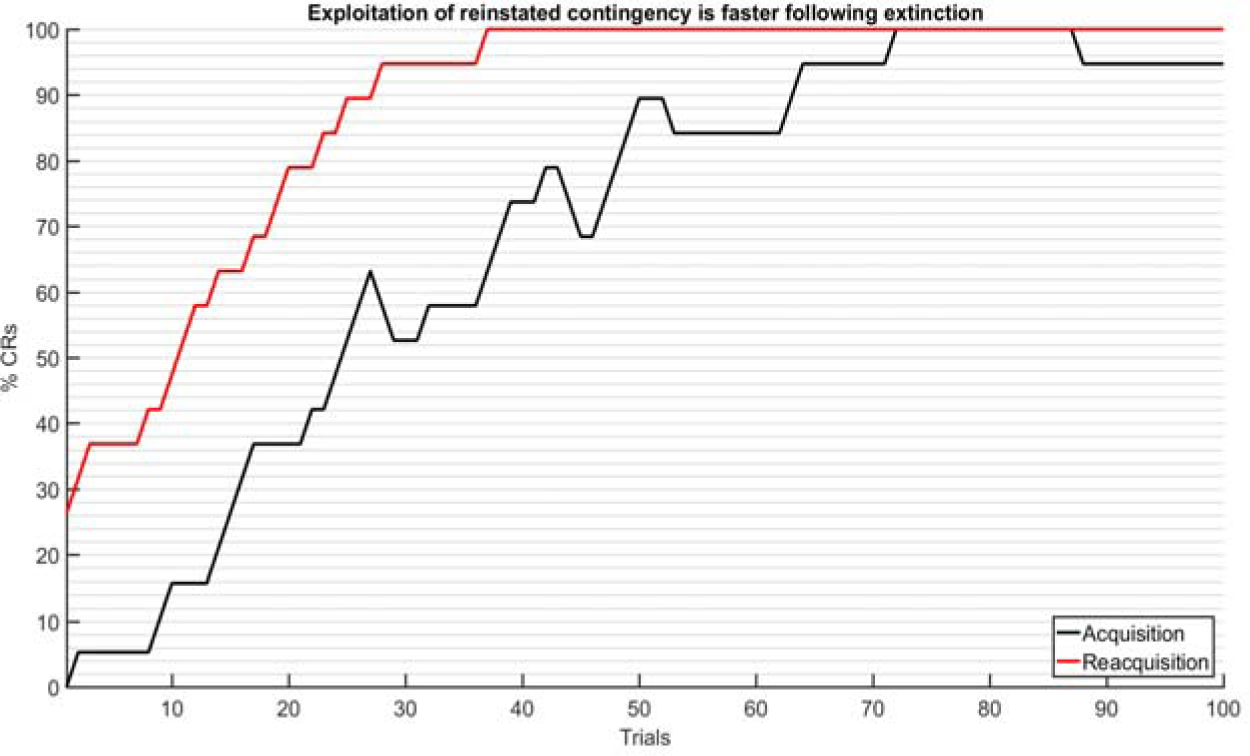
“Savings” shown by faster reacquisition. Superposing the acquisition and reacquisition phase results from Figure 9a on the same time line shows that the model’s post-extinction reacquisition of a conditioned response (red line) is faster than the initial acquisition of that conditioned response (black line), indicating that memories (here, weight increments) formed by the initial learning were “saved” despite the extinction training. The model’s faster reacquisition reproduces an often replicated experimental result. The property emerges even if initial acquisition has not been prolonged enough to cause a transition of control between the secondary and primary compartments of the model.

**Figure 5c.**
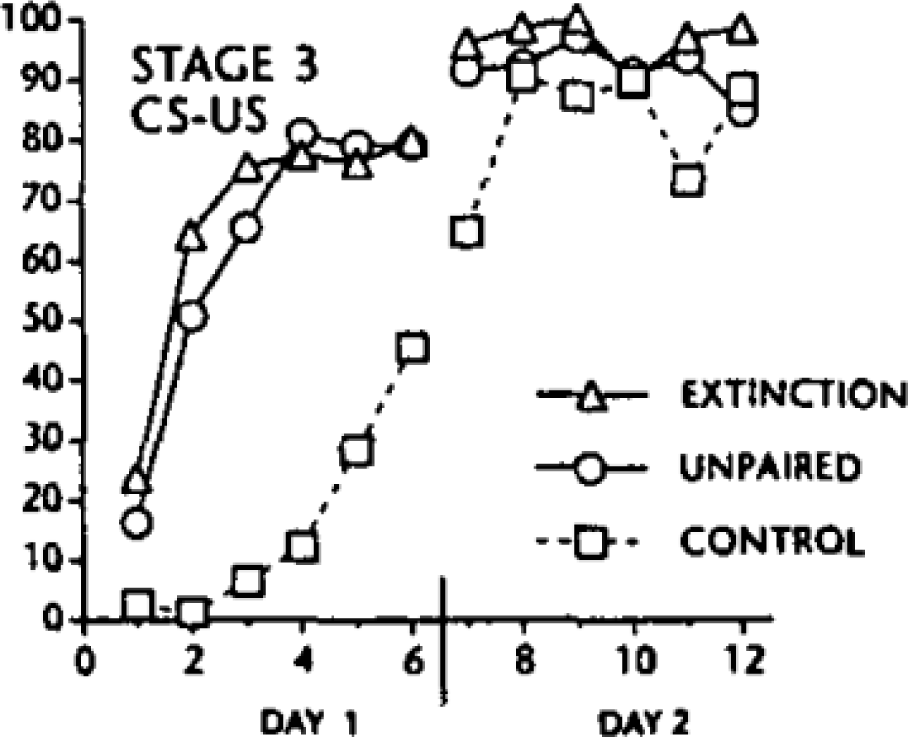
Figure reproduced from Kehoe & Macrae (1997). This figure shows rapid reacquisition of a Pavlovian conditioned response following extinction (solid line, triangle marker) alongside the initial acquisition of that response by a naïve control (dashed line, square marker), with results similar to those depicted in Figure 5b. Rapid reacquisition is also seen for cued instrumental responses.

The model’s rapid reacquisition results from the parallel contributions of the GO and NOGO pathways. The synaptic weights of the NOGO pathway are adjusted opposite to the GO pathway, and undergo larger reinforcement-related adjustments than those of the GO pathway. Compared to the naive state, in which the weights of the GO and NOGO pathways are roughly equal, the extinction-induced weights of the GO and NOGO pathways are lopsided in the favor of the NOGO pathway (Figure 5d), as a result of its faster learning rate. When a reward contingency is re-established following extinction, learning re-facilitates the rewarded response via both reengagement of the GO pathway and by disengagement of the NOGO pathway. Compared to initial acquisition of a rewarded response, its reacquisition is accelerated by rapid disengagement of the NOGO pathway, which “dis-occludes” already elevated associative links to the reengaging GO pathway.

**Figure 5d.**
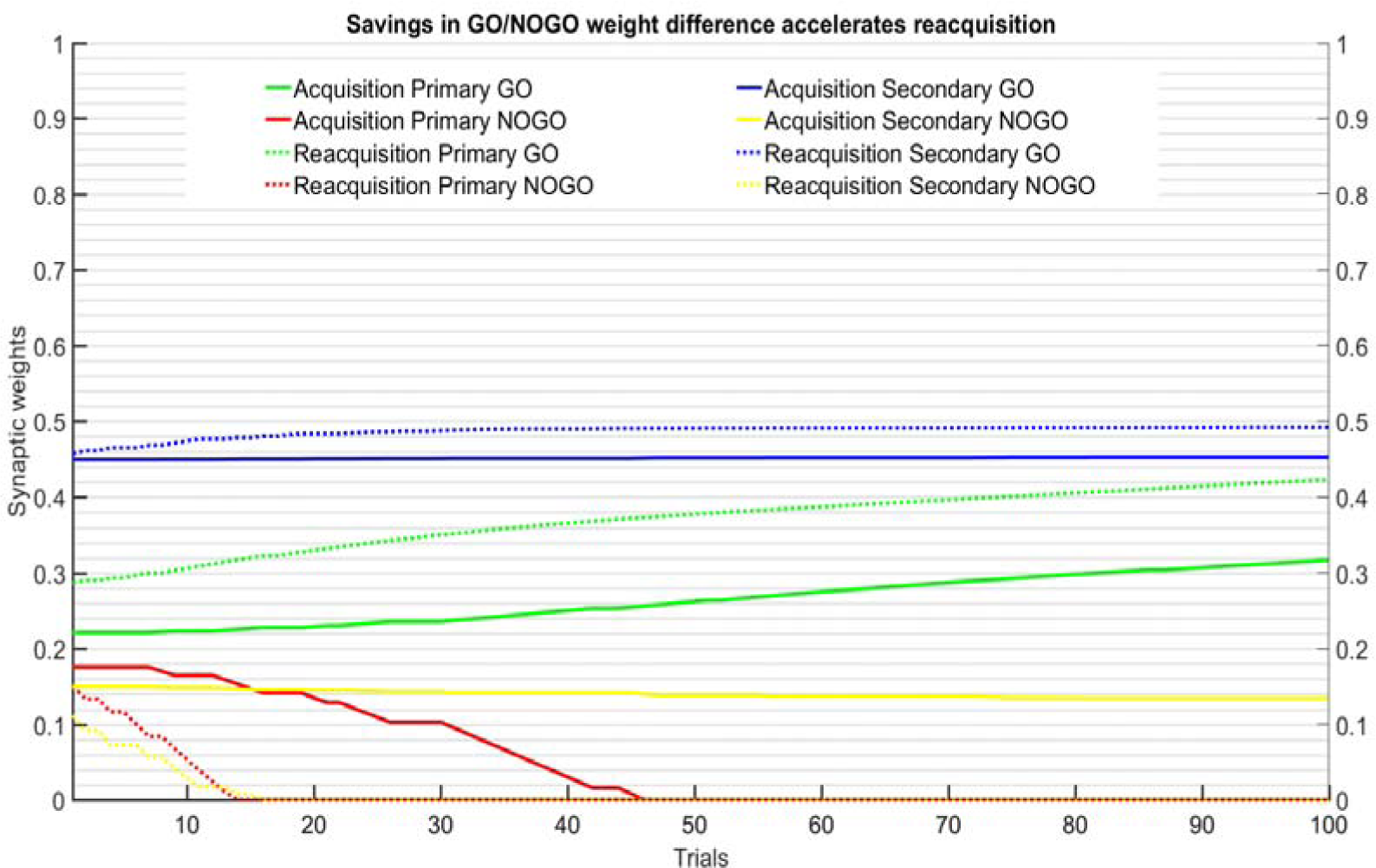
Model synaptic weights during initial acquisition and reacquisition. This figure depicts the initial values and changes over time of synaptic weights in both the GO pathway (corresponding to synapses onto striatal D1-type MSPNs, which promote behavior; green and blue lines) and the NOGO pathway (corresponding to synapses onto striatal D2-type MSPNs, which oppose behavior; red and yellow lines). At the onset of the reacquisition block (dotted lines), GO weights are slightly elevated relative to the beginning of the initial acquisition (solid lines) while NOGO weights are lowered. The difference between the GO and NOGO pathways resulting from extinction, combined with rapid disengagement of the NOGO pathway during reacquisition and engagement of the rapid-learning secondary compartment (blue and yellow lines) during reacquisition, contributes to the savings effect observed in Figure 5b.

Figure 6 shows model performance for a reversal learning paradigm, in which the contingency linking a stimulus to a rewarded response repeatedly switches between two alternative responses. This figure demonstrates the repeatability of control transfers between primary and secondary compartments, as well as a savings effect for both accuracy and habitual (model-free, DLS) control. The three traces shown in Figure 6 correspond to three separate reversal phases in chronological order, from top to bottom. The color of each point composing the solid traces codes the compartment in control of behavior: either the primary compartment (red) or secondary compartment (blue).

**Figure 6.**
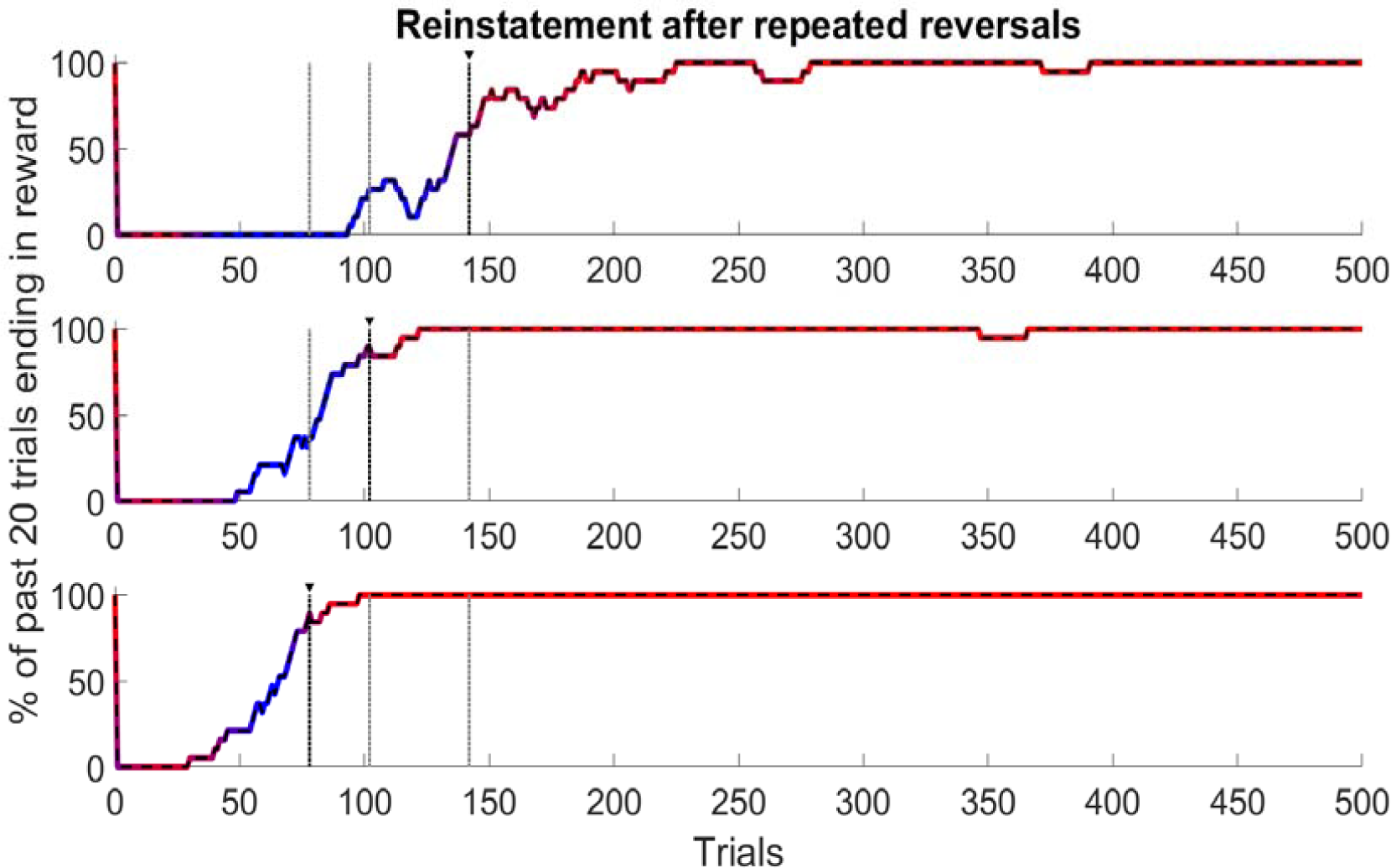
Reversal learning and habit formation facilitated by repetition. A 20-trial moving average percentage of correct (rewarded) responses is depicted across a series of rule reversals from top to bottom, with each graph starting at the change of reward contingency from a 500 trial period rewarding stimulus 1-response 1 (not shown) to 500 trials rewarding stimulus 1-response 2 (trials 0-500 shown in figure), over a series of three repetitions of this reversal paradigm for a total of 3000 trials. The color of each point composing the solid traces codes the compartment in control of behavior: either the primary compartment (red) or secondary compartment (blue). A black vertical dotted line beneath a black triangle indicates the first stable transition to primary compartment (habitual) control (i.e., when the model DLS first makes 90% of the decisions in a 20-trial span), whereas gray vertical dotted lines are included to provide a comparison with the transition points in the other trial blocks. Over repeated rule reversals, the model’s accuracy improves within fewer trials (compare solid red-blue traces), and the transition to primary control occurs within fewer trials (compare vertical dotted lines).

Across repeated reversals, two principal effects are visible: the model recovers correct performance more quickly following later reversals (a finding that extends the savings effect observed after one extinction as depicted in Figure 5) and the number of trials required to reinstate primary compartment (DLS) control of behavior decreases with each reversal. The model thus exhibits a progressive tendency to “relapse” to a previously-extinguished habit following restoration of favorable outcomes.

Whereas the savings effect shown in Figure 5 is dependent on relative differences in the rates of learning in the GO and NOGO pathways, the relapse effect shown in Figure 6 arises from differences in both learning rates and weight saturation limits between the primary and secondary compartments. Following each reversal, reinforcement training occurs beyond the training level at which the secondary compartment’s weights reached their saturation limit, whereas the primary compartment’s weights continue to be incremented. This builds a bias in favor of primary compartment control as trials accumulate. *During* each reversal, the GO weights of both primary and secondary compartments (for the now non-reinforced response) are decremented, but the rapid increments of the NOGO weights results in a demotion of the non-reinforced behavior long before GO weights in either compartment are decremented to values seen before the most recent block of reinforced trials. The net result is a gradual accrual of bias in favor of the primary compartment over multiple reversals, and faster habit-reinstatement across successive reversals, consistent with empirical reports (e.g., Clarke et al., 2011) on dopamine-dependent reversal learning.

Figure 7 shows the behavior of the model in an experimental paradigm that mimics the plus-maze paradigm shown in Figure 1. The experiment consists of three trial blocks. The first is a random exploration block corresponding to exploration of the plus maze from a random starting position. During the first trial block, each of two secondary representations, E and W, are linked by reinforcement or non-reinforcement to responses 1 (approach) or 2 (avoid), respectively. These representations, which have an effect on only the secondary compartment, simulate the spatial codes by which a learner can orient itself in the plus maze (with each corresponding to a distinct maze arm). Thus, the secondary compartment is construed as having access to a sense of spatial orientation that is not directly accessible to the primary compartment. The second trial block consists of repeated presentations of stimuli that evoke the secondary representations in parallel with an independent primary stimulus representation, J, construed as merely coding a maze junction. This block simulates reinforcement training over repeated insertions into the same arm of the plus maze, during which a link between detecting J and turning right or left can be established. The final trial block consists of a switch in the insertion point, and activations of secondary (E, W) and primary (J) representations.

**Figure 7.**
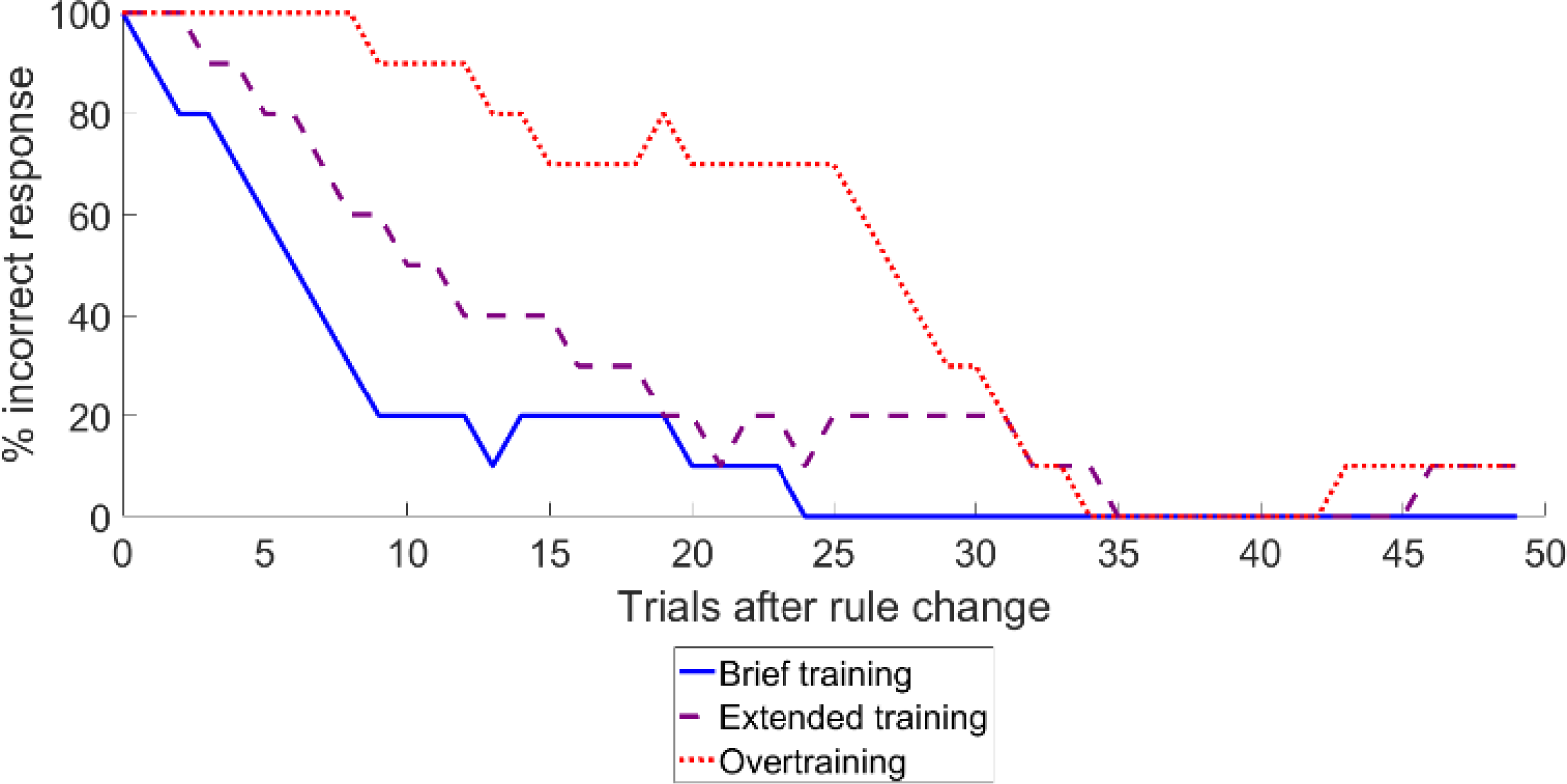
Effect of training period on incorrect response perseveration. A 10-trial moving average is presented of the percentage of incorrect responses produced after a change in reward contingency in which stimuli distinguishable within the secondary compartment, but not the primary compartment, cue a change in rewarded response. Longer periods of training correspond to more complete transfers of control to the primary compartment and, as a result, are associated with longer periods of incorrect response perseveration. The number of training trials per case plotted was: brief, 100 trials; extended, 1000 trials; overtraining, 2000 trials.

The behavior of the model during the third trial block depends on which compartment controls behavior at the end of the second trial block. The primary compartment’s recommended response will remain the same from the end of trial block two to the start of three, and would yield an incorrect decision early in block three, just after the switched insertion. Whether behavior is controlled by the primary or secondary compartment at the onset of the third block is dependent on the number of repeated reinforcements during the second block, because a sufficiently extended training period results in transfer of control to the primary compartment. The model’s incorrect behavior perseverates as long as the primary compartment remains in control of behavior, with acquisition of reward only occurring after the secondary compartment regains control following adjustment of the primary compartment weights by repeated failure to obtain reward. These findings are consistent with those of Smith & Graybiel (2013), in which overtraining was linked to both devaluation insensitivity (not modeled here) and increased activity in the DLS. Although extending the model to simulate how devaluation insensitivity increases with prolonged reinforcement training (Williams, 1938) is beyond the scope of this report, the model reveals why the degree of transfer to DLS varies directly with the extent of overtraining.

Figure 8 shows the response onset latencies, or reaction times (RTs), of the model over repeated reinforcement trials with two stimuli, each presented in a block for 1500 trials. In the first trial block, a stimulus that is codable as reward-predicting, by cortical representations that send afferents to both the primary and secondary compartments, is presented while one response choice is consistently rewarded. In the second trial block, a distinct stimulus, which is reward-predicting as coded by cortical representations that send afferents to the secondary but not to the primary compartment, is presented while one response choice is consistently rewarded. In both trial blocks, the time taken to decide on a response steadily decreases over the course of reinforcement. However, transfer of control to the primary compartment only occurs in the first block, because only in that block does that compartment receives afferents from a representation that distinguishes the reward-predicting aspect of the stimulus. Control transfer results in markedly lower RTs, because the primary compartment’s higher synaptic weight results in faster integration of afferent signals at the decision stage. In the second trial block, no such transfer occurs, and the speed gain due to practice is smaller, due to the smaller asymptotic synaptic weights achievable in the secondary compartment.

**Figure 8.**
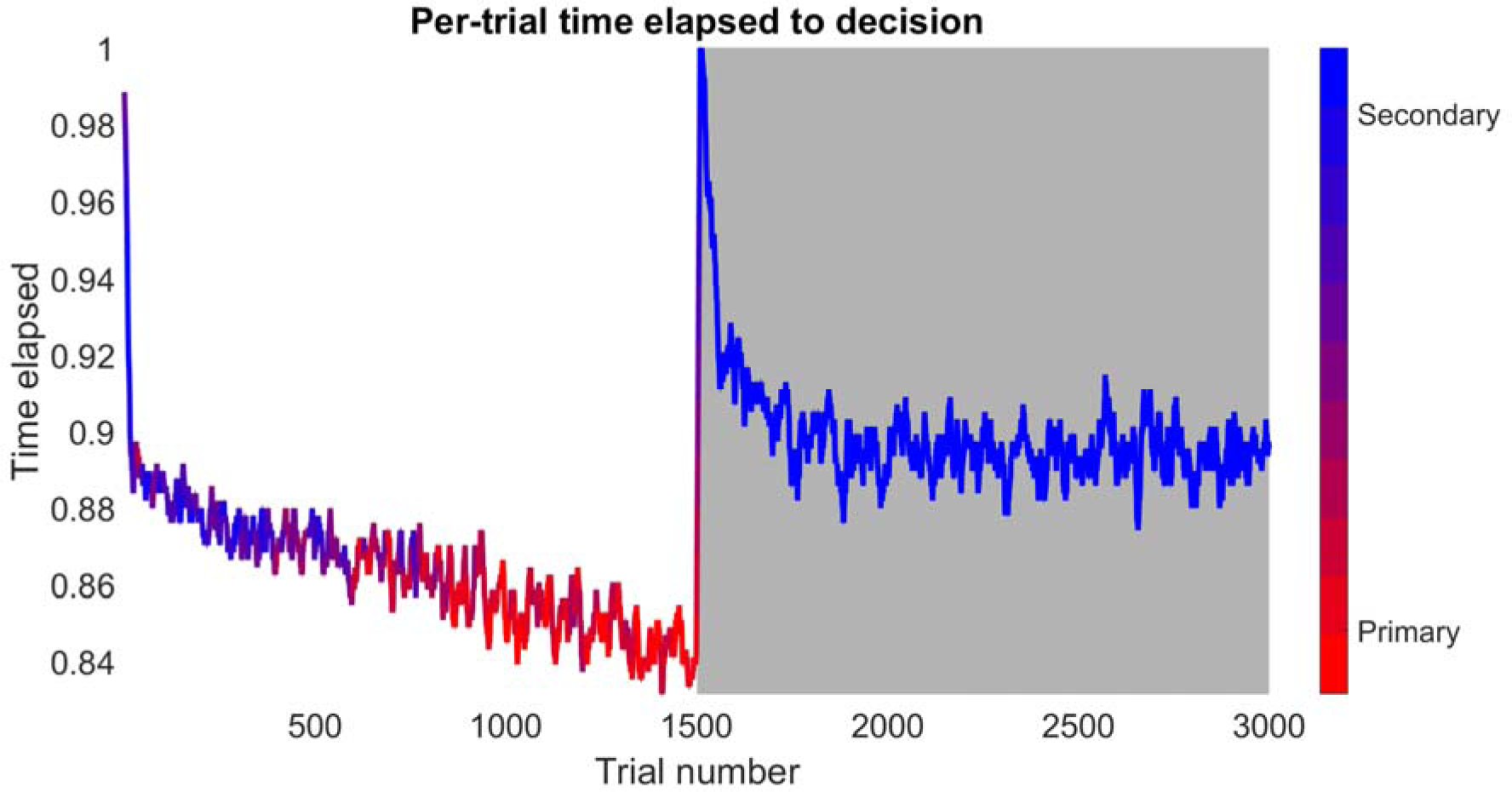
Evolution of reaction times over the course of reinforcement training. The reaction time needed depends on which compartment is in control of behavior, which itself depends on whether the compartment has access to a reward-predictive representation. The primary compartment’s higher asymptotic weights makes it capable of faster decisions after many trials, but transfer of control to the primary compartment – shown by a trace changing color from blue to red around trial 500 – only occurs if the stimulus being presented is reward-predicting as coded by afferents projecting to the primary compartment. After trial 1500, presentation of a new stimulus, which is reward-predicting as coded by afferents to the secondary but not the primary compartment, does not support a transfer of behavioral control – the trace remains blue – and so the asymptotic reaction time is longer. The time elapsed (y axis) from stimulus onset to response selection is normalized in order to depict the relative impact of training, which reduces the time taken for each decision to ∼90% of baseline in the secondary compartment and ∼80% in the primary compartment. The speed gained by practice is larger in the primary compartment due to its larger asymptotic weights.

Figure 9 shows the simulated effect of a DAT antagonist such as cocaine (Carboni et al., 1989; Budygin et al. 2002; Phillips et al., 2003; Budygin et al. 2007) on habit formation. Cocaine has an observed asymmetry of effect between ventral and dorsal zones of the striatum (Takahashi et al., 2008; Takahashi et al, 2009), particularly in the context of extinction (Fuchs et al, 2006), and is known to interact with dopamine transporter (Izenwasser et al., 1990; Guindalini et al., 2006) as well as dopamine receptors (Noble et al., 1993; Noble, 2000) and other cellular mechanisms (Venton et al., 2006). Cocaine’s DAT antagonism, which allows reward-induced DA elevations to persist longer than usual, is simulated as an elevation of learning rates in both compartments. To reflect the higher natural DAT level in DLS that is subject to such antagonism, the learning rate elevation (due to DAT antagonism) is greater in the primary (50% increase) than in the secondary compartment (20% increase). The larger learning rate gain in the model’s DLS leads to earlier and stronger engagement of the primary compartment, hence a more rapid transition to habitual control.

**Figure 9.**
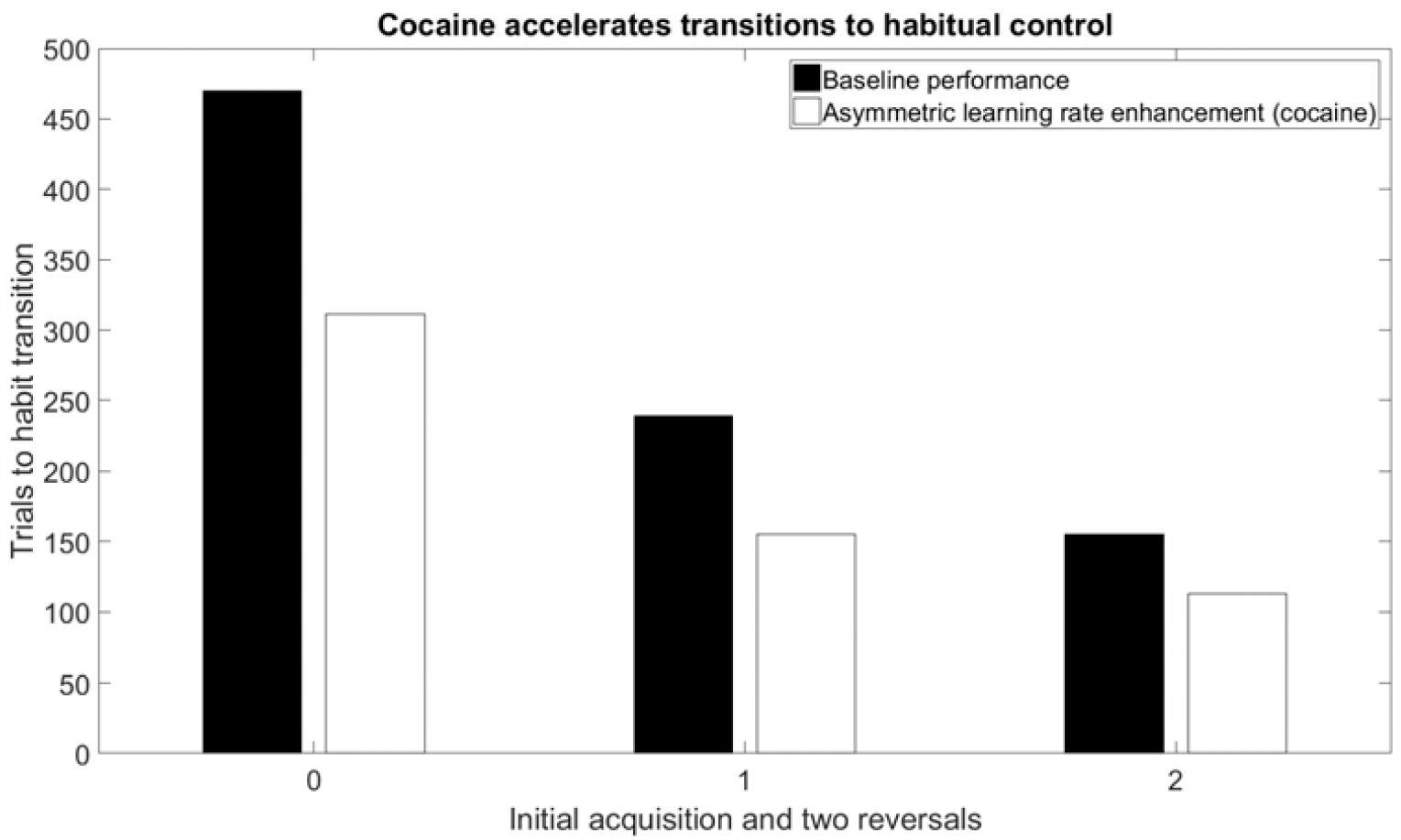
Simulating a DAT antagonist accelerates transition of control to the primary compartment. Under baseline parameterization (black), repeated reinforcement of a response leads to transition to exclusive control by the primary compartment that becomes faster following repeated reversals (cf. Figure 6). An alternate parameterization corresponding to a DAT antagonist, such as cocaine (white), leads to a faster transition of control to the primary compartment, both during the initial acquisition and following repeated reversals. Data shown correspond to initial acquistion and two reversals occurring on trials 1-500, 1001-1600, and 2001-2500, respectively; only those blocks corresponding to a single (i.e., non-reversed) contingency are shown.

Figure 10 demonstrates a tradeoff between speed and accuracy under conditions in which the secondary compartment is capable of performing to a higher degree of accuracy than the primary compartment. In this simulation, one of three responses is rewarded, contingent on the presentation of a random combination of primary and secondary stimuli. The contingencies linking stimuli to reward were chosen to ensure that prediction of reward based on primary stimuli alone can achieve no more than 50% accuracy, whereas inclusion of secondary stimulus information allows for perfect accuracy.

**Figure 10.**
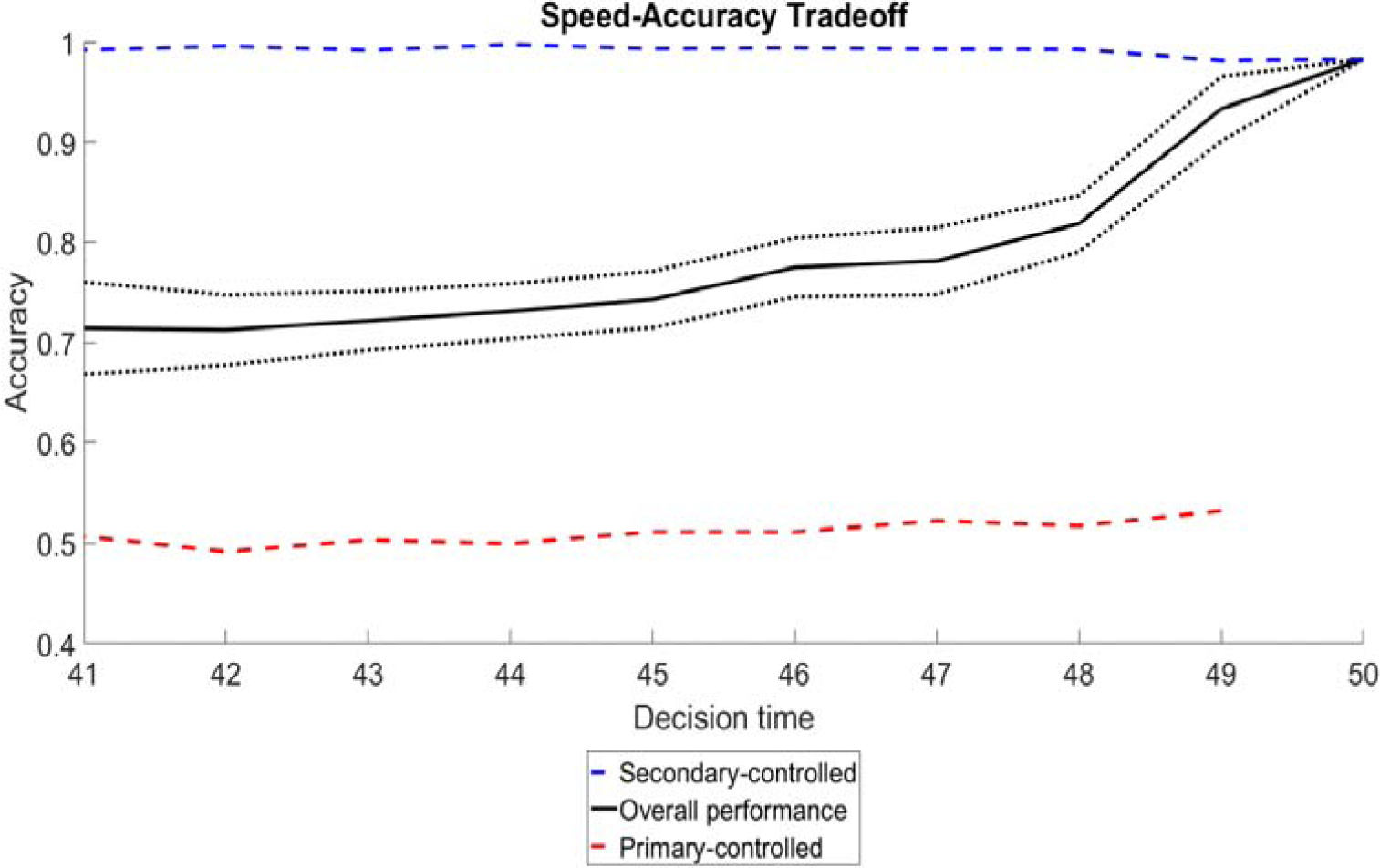
Tradeoff between speed and accuracy as a result of competition between primary and secondary compartment. Trials are performed under a paradigm in which the primary compartment is capable of achieving a maximum of 50% accuracy (red dotted line), whereas the secondary compartment can achieve 100% accuracy (blue dotted line). Both compartments are capable of producing a response within the window of 41 to 50 time steps; however, as the time taken to produce a response increases, the proportion of responses driven by the secondary compartment increases, improving the mean accuracy of the model’s performance (solid black line). The figure is based on 2000 trials, and the dotted black line indicates the standard error of the mean accuracy for each bin of decision times.

For the Figure 10 simulation, the increase in accuracy is solely a consequence of a larger proportion of decisions being controlled by the secondary compartment, which requires longer decision times to accommodate the longer stimulus-processing delay associated with computing the more-predictive representations that send afferents to the secondary compartment. This figure reproduces the longstanding experimental observation of a speed-accuracy tradeoff (SAT) in forced choice paradigms (Pew, 1969). Explaining this SAT as a result of a mixture of lower-quality decisions by the primary compartment and higher-quality decisions by the secondary compartment agrees with prior modeling results suggesting that other SATs may be due to a competition between parallel strategic processes (Maanen, L. 2016; Keramati et al. 2011; Boldini et al. 2004).

**To summarize, the simulation results shown so far flow mainly from two sets of model parameters – learning rates and weight asymptotes – that were motivated by neurobiological findings listed in Table 2. Table 2 highlights these key parameters, along with three ancillary factors, and their behavioral effects. These attributions were tested by examining deviant parameter choices. For example, let the primary and secondary weight limits be denoted by Γ_P_ and Γ_S_ respectively, and the primary and secondary learning rates by λ_P_ and λ_S_. All prior simulations were performed with the values: Γ_P_ = 1; Γ_S_ = 0.3; λ_P_ = 0.003; λ_S_ = 0.0045. These parameter choices, while informed by biological constraints, represent a single sample of the space of possible parameterizations. If absolute values of parameters of a given type are not too dissimilar, then what matters most are the rank orderings of the values. This can be seen in**

**Table 2.** Choice of model parameters. Four design choices central to the model’s functionality were motivated by anatomical evidence.

| Model parameter | Behavioral effect | Supporting evidence |
| --- | --- | --- |
| Faster learning rate in secondary compartment | Secondary compartment maintains control of behavior early in training | Hurd et al. (1994); Gonzales-Hernandez et al. (2004); Collier et al. (2011); Calipari et al. (2012) |
| Higher learning asymptote in primary compartment | Primary compartment gains control over behavior after repeated reinforcement | Mancuso et al. (2014); Meredith et al. (2008) |
| Higher baseline weight values in secondary compartment | Secondary compartment controls behavior in the absence of reinforcement | Villalba & Smith (2013) |
| Longer persistence of reward-induced learning signals in primary than secondary compartment | Primary compartment gains control over behavior following periods of extended reinforcement | Willuhn et al. (2012); Ito et al. (2000, 2002) |
| Asymmetrical crossover of learning signal from secondary to primary compartment | Prevents secondary compartment from locking out primary compartment | Haber et al. (2000) |

Figure 11, which reports results of swapping rankings, for one parameter type at a time. For all cases summarized in Figure 11, the model was rewarded at p=1.0 for one response to explore the effects of the four learning parameters on weight trajectories and behavioral switching. With rate parameters λ_P_ and λ_**S**_ swapped, initial control by the secondary compartment is preserved as long as its weights are initialized to significantly higher values than primary compartment weights; however, the crossing point of the secondary and primary weight trajectories occurs much earlier in training, resulting in more rapid transition of control to the primary compartment. In contrast, exchanging the weight asymptote parameters Γ_P_ and Γ_S_ alone, without also making λ_**P**_ much greater than λ_S_, prevents any transition of control from occurring. There is no crossing point between the weight trajectories.

**Figure 11.**
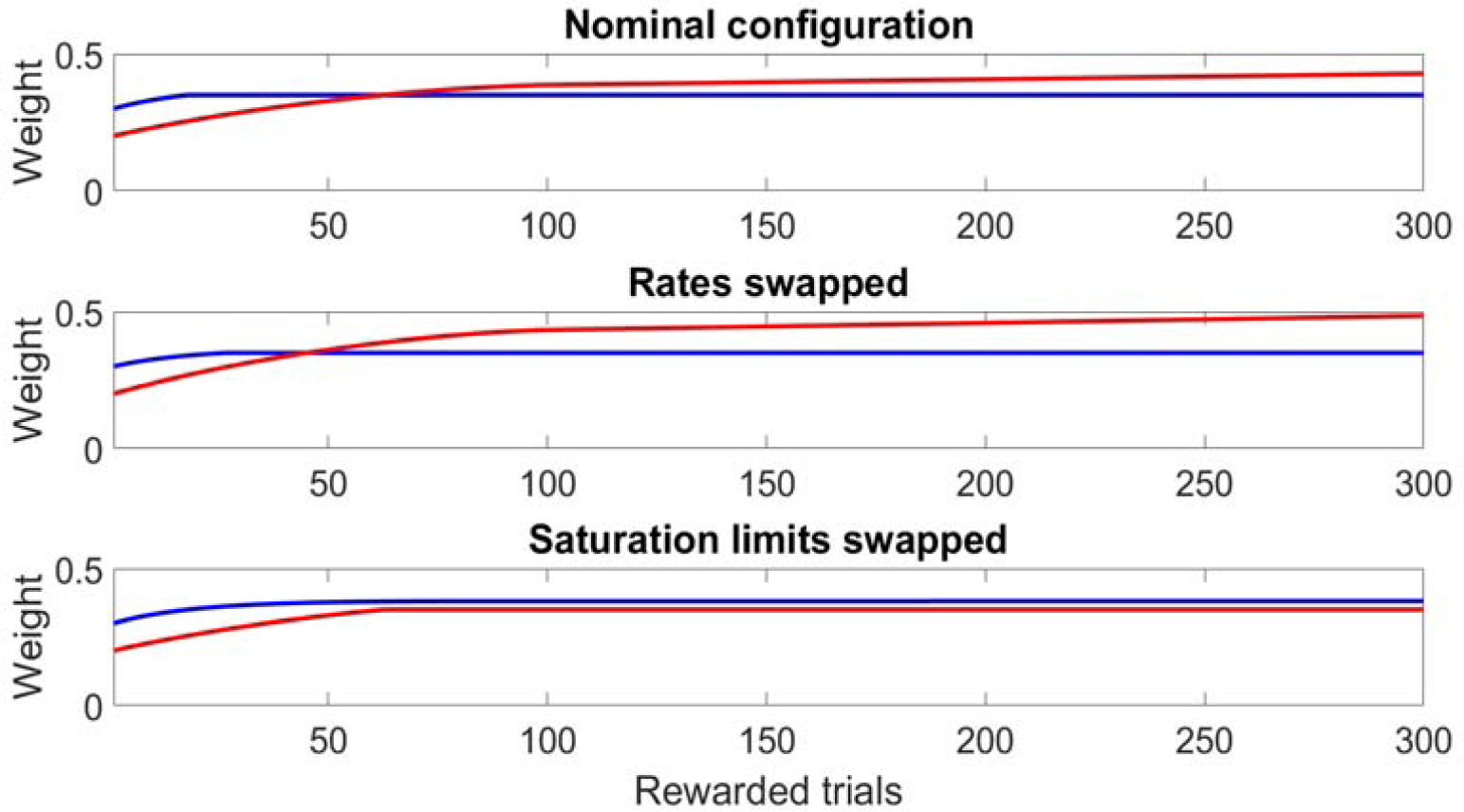
Effects of changing the rank orderings of learning parameters across two compartments. This figure depicts weight trajectories in both the primary and secondary compartments as rewarded trials accumulate, the scenario previously depicted in Figures 1 and 7. Each compartment has two parameters subject to variation, the learning rate *λ* and weight saturation bound, Γ. The topmost graph in this figure depicts the standard configuration of the model, with *λ* = 0.0045 and Γ = 0.35 in the secondary compartment and *λ* = 0.0036 and Γ = 1 in the primary compartment. In the middle graph, the learning rates have been swapped between the two compartments, such that *λ* = 0.0036 in the secondary compartment and *λ* = 0.0045 in the primary compartment. This results in an earlier crossing point from secondary to primary control, reminiscent of the simulated cocaine parametrization shown in Figure 10. The bottom graph shows a parameterization in which the saturation limits were Γ = 0.35 in the primary compartment and Γ = 1 in the secondary compartment, resulting in a system in which the primary compartment weight never exceeds that of the secondary compartment – a pathological state implying an inability to enter habitual mode.

A major factor in the habit-forming potential of a response contingency is the predictability with which it leads to a reward, because burst releases of dopamine, which gate weight changes, reflect reward prediction error (RPE), which decreases as a reward becomes more predictable (Schultz, 1999). The DMS exhibits a stronger decrease in prediction-related dopamine release than the DLS. This physiological asymmetry is represented computationally within the model as differences in the drop-off of weight adjustment magnitudes as a function of reward predictability, with the secondary compartment having a larger dropoff (95% lower magnitude) compared to the primary compartment (50% lower magnitude) at maximum predictability. Due to these differences in prediction-related learning changes between compartments, variations in the predictability of reward will affect the transition to habit mode. Lower predictability is expected to favor the secondary compartment in the bid for control by preventing the comparatively large prediction-related decreases in weight adjustment from occurring. Drops in predictability of reward are expected to result not only in slower acquisitions (due to the lower density of trials that end in a reward), but also in a slower transition from secondary to primary control of behavior, as a result of the asymmetry in predictability-related effects on learning between the two compartments.

The repeated reversals paradigm depicted in Figure 6 was revisited in new simulations, but this time with less than perfect predictability of reward. **Figure 12** shows the simulation results for a case in which only a random 75% of correct responses lead to a reward, while the remaining 25% of correct responses result in non-reward, as do all incorrect responses. As expected, under this probabilistic regime both the transition to habitual control and the re-acquisition of the rewarded response are delayed, relative to the p=1 result shown in Fig. 11.

**Figure 12.**
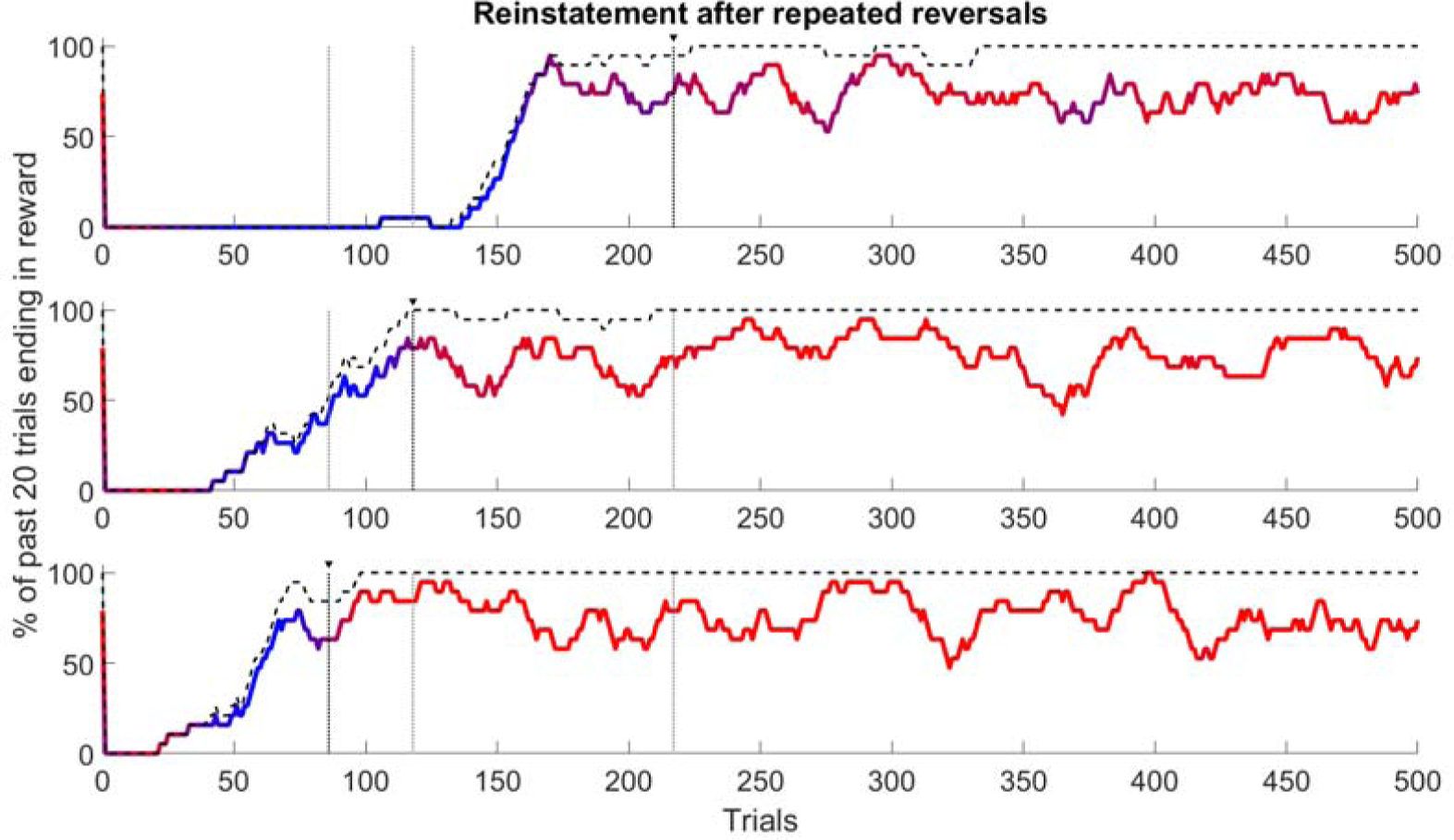
Behavior of model under a probabilistic reward schedule. In this case the probability that the rewarded response will lead to a reward is p=0.75 (compared to p=1 in the baseline case.) The black dashed trace running above the colored trace represents the percent correct responses, i.e., the percentage of responses that would be rewarded if p=1. Although the model still reaches peak accuracy (∼ 75% of responses achieve reward), both the initial acquisition of the response and the transition to primary compartment control are delayed in comparison to the p=1 case. To facilitate comparisons across repeated acquisitions, each panel has dotted vertical lines corresponding to times of stable transition to primary compartment control. One of the three dotted verticals, namely the darkest one marked by a superposed black triangle, corresponds to the trial block plotted in that panel. The two fainter dotted verticals mark the transition points from the other two panels. The transition time occurs earlier with each successive reversal, but always later than if p=1 (compare Fig. 6).

**Figure 12** shows that in the model, the introduction of uncertainty to a contingency delays the shift to habitual control. This fits with the proposal of Daw et al. (2005) that an adaptive system should prefer model-based control to model-free control under conditions of uncertainty. This delay in switching, and all control switching effects presented so far, resulted entirely from slow, learning-related changes to synaptic efficacy that in turn depend on phasic RPE-related dopamine release events. Are there other effects of DA that, if included in the model, would work to delay the shift to habit under conditions of uncertainty? One reason to think so is that *in vivo*, DA also has more immediate effects, on the time-scale of a single decision. Notably, the instantaneous level of DA controls the balance between the direct and indirect pathways, because it facilitates the direct pathway via D1Rs and dis-facilitates the indirect pathway via D2Rs (review in Gerfen & Surmeier, 2011). This can affect performance vigor and rate, as attested by the street name, “speed”, for the class of DA agonists known as amphetamines. Such effects are entirely consistent with the current model, although how much a DA agonist can change the direct-indirect balance depends on the history of learning, individual or regional differences in DAT expression, etc.

To exemplify this issue, a simulation was run to address the effect of the so-called “uncertainty” or “risk” response of dopamine cells (Fiorillo et al., 2003; Tan & Bullock, 2008a, 2008b; Schultz, 2015), which is seen only on probabilistic, i.e., uncertain, reward schedules. The “uncertainty” response is an above-baseline, slow, DA-release event, not a burst DA-release event. For that reason, it can be expected to have a much larger effect in low-DAT regions of striatum than in the high-DAT DLS: in the DLS, the slowly released DA could be cleared almost as fast as it was released. To model single-trial facilitation due to DA release, a parameter β ≥ 0 was introduced with separate values, β_secondary_ and β_primary_, in the two compartments. A larger β corresponds to a stronger transient facilitation of the direct, and dis-facilitation of the indirect, pathways in the model striatum. The effects of β are shown in Figure 13, from a simulation in which β was non-zero only in the secondary compartment, and in Figure 14, in which β was nonzero in both compartments, but larger in the secondary compartment. Both cases are consistent with the expectation, and general model assumption, that lower DAT levels in the DMS result in a greater single-trial effect from DA release events than a comparable release in the DLS (regardless of whether the putative mechanism for such an effect is related to phasic, learning-related DA effects, as set by the λ learning rate parameter, or due to performance-related DA effects, as set by the β parameter.) Figure 13 illustrates that the uncertainty response of DA cells can indeed delay the transition to habitual control if DAT levels are such that the DA signal has a significant effect in the DMS but a negligible effect in the DLS. Figure 14 shows an intermediate case between **Figure 12** and 13, in which an uncertainty response DA release that is not fully negated by DAT in DLS does little to delay the transition to habit by the time of the third reversal.

**Figure 13.**
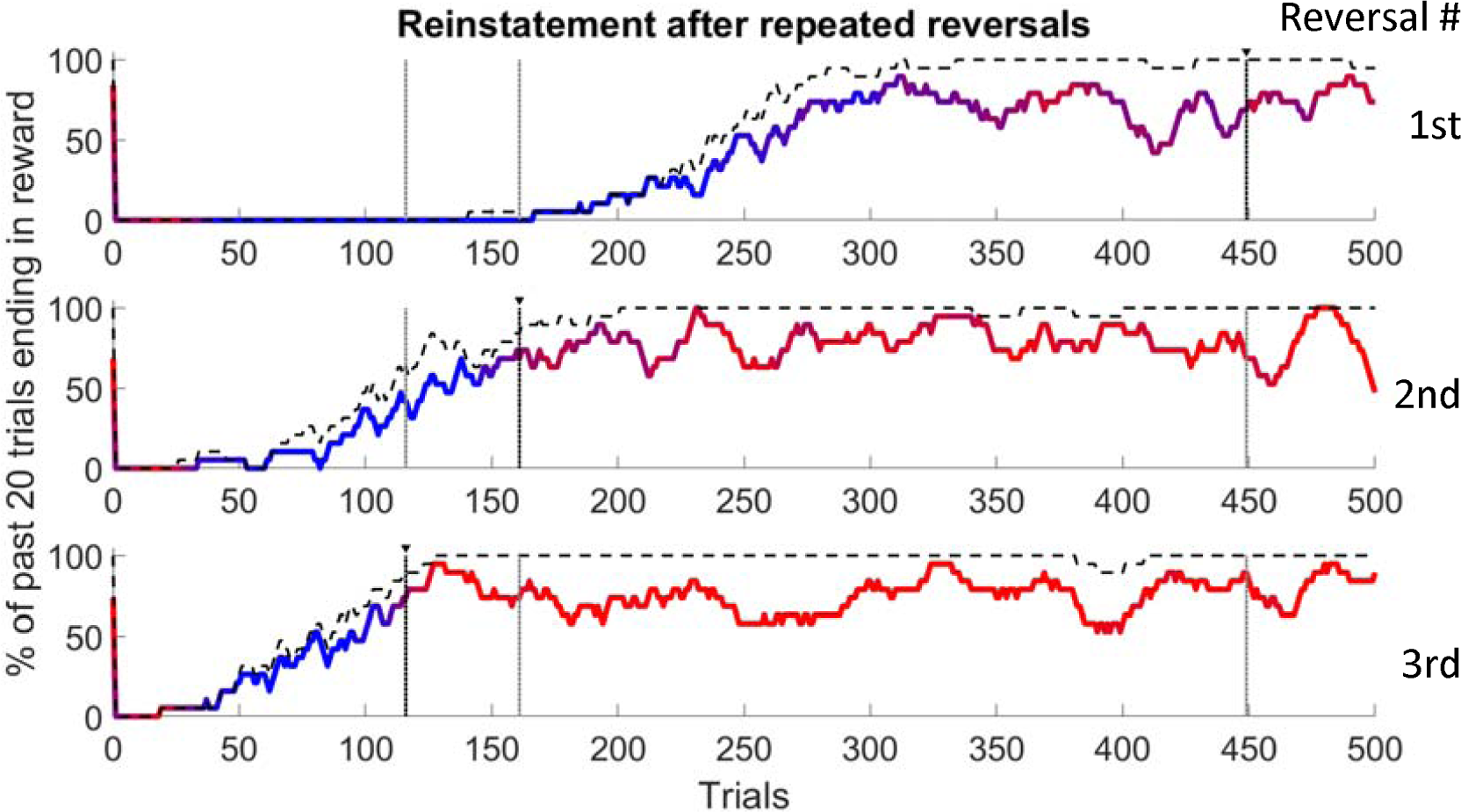
Transitions when dopaminergic uncertainty responses affect only the secondary compartment. The model was configured to include a nonzero β parameter in the secondary compartment only (β_secondary_ = 0.2, β_primary_ = 0.0) to simulate an immediate effect of dopamine not mediated by a persistent weight adjustment. As in Figure 12, the transition from secondary compartment control (blue) to primary control (red) is delayed compared to baseline, as is reacquisition of the active contingency following reversal. Non-learning impacts of dopamine on the decision making process, as represented by the β parameter, may thus be sufficient to delay the transition to habitual decision making in addition to the learning-related effects of predictability shown in Fig. 12. As in Fig. 12, the dotted vertical lines correspond to the earliest stable transition to primary compartment control, with the black triangle over a black dotted line corresponding to the current trial block while the gray dotted lines show the transition times within the other two trial blocks for comparison.

**Figure 14.**
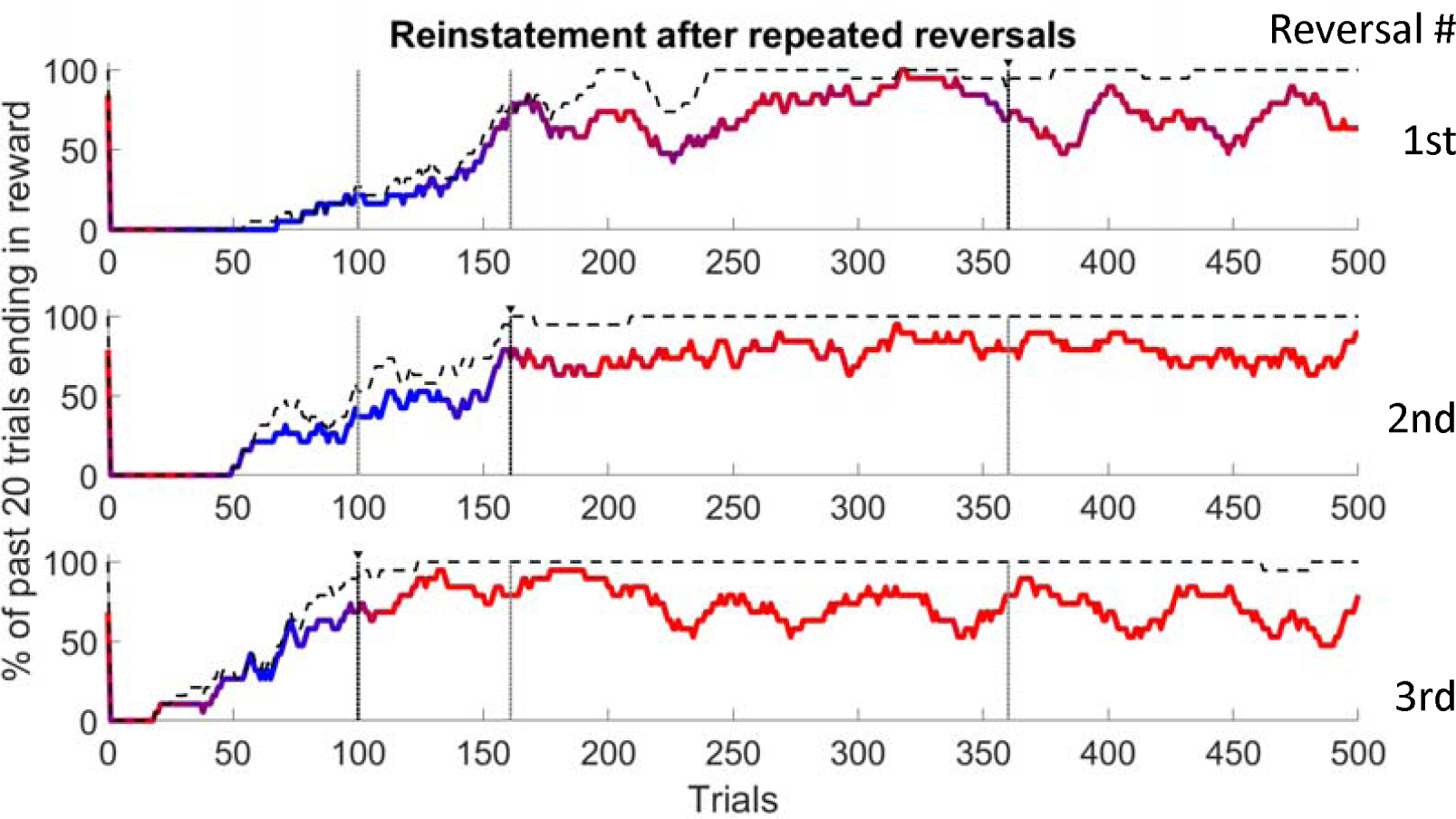
Transitions when dopaminergic uncertainty responses affect both compartments. Behavior of model configured to include a nonzero β parameter in both compartments, with a smaller β in the primary compartment (β_secondary_ = 0.2, β_primary_ = 0.1) simulating conditions in which non-learning effects of DA are elevated in both subregions of striatum. As in Figure 13, the transition from secondary compartment control (blue) to primary control (red) is delayed compared to baseline, as was reacquisition of the response following reversal. In contrast to results shown in Fig. 13, the more rapid transition to primary control suggests that the balance of short-term dopamine effects between sub-regions of striatum may be an important factor in habit formation under probabilistic contingencies, and that non-learning effects of DA, as represented by the β parameter, may counter the effects of prediction-related changes in dopamine-mediated learning. As in Fig. 12, the dotted vertical lines correspond to the earliest stable transition to primary compartment control, with the black triangle over a black dotted line corresponding to the current trial block while the gray dotted lines show the transition times within the other two trial blocks for comparison.

## Discussion

The model simulated here assumes that reward-related signals projected by midbrain dopamine neurons to two distinct striatal sectors – DMS and DLS – interact with sector-specific parameters that control the rates, and asymptotes reached, of changes to the synaptic efficacies of fibers (e.g., from cortex) that can mediate cued excitation of the medium spiny projection neurons (MSPNs) that give rise to the direct (action promoting) and indirect (action demoting) pathways through the basal ganglia. The simulations show that these interactive processes are sufficient to both enable and mediate reward-guided transfers of behavioral control. Characteristics of the transfers produced during the simulations conform to characteristics of analogous *in vivo* transfers.

Notably, initial acquisition of rewarded actions involves fast learning and early control by DMS, whereas over-training (well past the point when choices are accurate) is needed to cause the transition to control by the slower-learning DLS. Moreover, extinction as well as uncertainty promote a return to control by the model’s DMS sector. Furthermore, reacquisition following extinction is faster than initial acquisition, and, relatedly, the number of training trials needed to reacquire a response after a contingency reversal declines across repeated experiences with the reversal. Because internal representations (of cues/contexts) with imperfect predictive validities are processed in parallel and race for control of stochastic decisions to launch responses, the decisions made by the model generate a mixture of reaction times, and these exhibit the well-established speed-accuracy tradeoff (SAT), provided only that representations with greater predictive validity take longer to compute than those with less validity at predicting the response that can earn reward.

In the empirical studies that have reported transfers of control, the focus has been on non-overlapping phases, and it seems that a common assumption is that only the DMS or the DLS would be in control at a given phase of learning. What our simulations reveal, for the first time, is that in a stochastic model that makes natural assumptions about parallel processing and incremental learning, there can be long phases during which control is very gradually passed from DMS to DLS, or the reverse. There can also be stable phases in which the degree of control is mixed, especially on probabilistic schedules of reward when the predictive validities of competing representations are similar. It is in such cases that the model can show the SAT simultaneously with a mixture of DMS- and DLS-mediated control within a block of trials.

To enable simple tests for the sufficiency of a few core mechanisms, the model omits many documented processes. For example, although training may produce persistent changes in neuron excitability that can bias action decisions in the absence of altered sensory-motor synapses, and that might affect transitions to habit in some species (Brembs, 2011), the model lacks any such process. However, by highlighting subcortical, reward-guided synaptic plasticity as a mechanism that is sufficient to switch behavior between multiple operating modes, the present study illuminates a huge range of typical cases.

Notably, the sufficient mechanism specified only uses circuitry believed to predate significant evolution of the pallium or cortex, which, beyond making the model applicable to many more species, also helps explain lesion data demonstrating that control transfers can survive various knockouts of cortical influence in mammals (e.g. Killcross & Coutureau, 2003). Yet, although the model described here does not require a reward-adapted “critic” in the cerebral cortex to accomplish transfers of control, the model is not incompatible with evidence that cortex provides ancillary bases for, and assists in (e.g., may accelerate), transfers of control among representations that affect action choices via striatal circuits. Prior models (e.g., Brown et al., 2004) have addressed data suggesting that DA-dependent plasticity in striatum synergizes with DA-dependent plasticity in frontal cortex to control decisions, and DA also modulates learned enhancements of task-controlling representations in sensory cortices (Bao et al., 2001; Holca-Lamarre et al., 2017). Cortical contributions to strategy shifts have been shown to be significant, especially in paradigms necessitating complex analyses of stimuli, selective attention to some stimulus features while ignoring others, and vigilant inhibition of prepotent (often habitual) behavior. Most such studies have been conducted in mammals (but see Krauzlis et al., 2017), and it remains to be seen whether the cortical contributions to control are themselves at least partly dependent on basal ganglia decisions, which exert control at multiple levels of the cortical hierarchy.

Recently, Thura & Cisek (2017) reported evidence from monkeys that cortical activity, but not pallidal activity, predicted the decision to reach for a target. This result might seem to contradict predictions of the present model, which relies (implicitly) on striato-pallidal pathways of the basal ganglia to guide decision making. However, the results of Thura & Cisek (2017) emerged within a unique paradigm, in which subjects began the experiment at a 50% criterion for each binary choice - that is, subjects were not conditioned to prefer one response to the other. Given such a training paradigm, cortico-striatal synaptic weights would equally favor both response options, so the model presented here predicts that activity at striato-pallidal stages would not provide a reliable indicator of the forthcoming decision. Instead, disparities in cortical activation between candidate responses would be solely responsible for decisions. Indeed, this often happens in the present model, which was made stochastic by using random noise to create learning-independent disparities at the cortical decision stage. This example further emphasizes the point that there is no reason to suppose that the sparse ancient mechanisms simulated here give a complete picture of decision making. They can coexist with many other factors (cortical, hormonal, etc.) in decision making.

Beyond being assumed as a source of context and cue representations that mediate model-based or model-free decisions, cortical contributions to decision making were deliberately excluded from the model to explore the competence of a minimal and ancient, but highly adaptive, circuit. Many studies support the truly ancient status of reward-responsive DA neurons, DA-dependent plasticity of decision-making, and a significant DAergic projection to a striatum, which is dominated by a single cell type, MSPNs, whose two major subtypes alternatively express either SP and D1Rs, or ENK and D2Rs (Reiner, 2009). These MSPNs give rise to direct and indirect pathways that course from striatum to pallidal and/or nigral territories. One other published treatment (Crossley et al, 2016) has simulated the ability to acquire, extinguish, and more quickly reacquire action with a sparse set of striatal resources that might be equally ancient. Although that study differed fundamentally from the present one – only one striatal sector, analogous to DLS, was modeled – it is instructive to note an abstract commonality, as well as key differences, between the models. The abstract point of agreement is that both models achieve savings by new learning in a pathway that can inhibit the expression of learning within a distinct, behavior-promoting, pathway.

This point of agreement extends as well to recent efforts to explain savings after extinction of acquired fear responses (e.g., John et al., 2016). However, rather than using plasticity of synapses onto MSPNs of the indirect pathway to demote actions during extinction, as in the present model, the model in Crossley et al. (2016) uses purported plasticity of synapses made by fibers from neurons in intralaminar thalamus onto cholinergic interneurons (ChINs) of the striatum. There are at least two problems with that model: first, it assumes that direct pathway MSPNs cannot fire unless ChINs pause their firing. However, recent *in vivo* and other data (reported and reviewed in Zucca et al., 2018) indicate just the opposite: ChIN pauses cause hyperpolarizations of, and reduced firing in, affected MSPNs. Second, although it might be possible to address the prior critique of a key assumption of Crossley et al. with another assumption, e.g., by reversing the assumed polarity – from LTD to LTP – of the extinction-induced plasticity at the synapses onto ChINs, there is a logical reason to prefer modeling extinction effects via plasticity of synapses onto indirect pathway MSPNs, rather than onto ChINs. Although ChINs may be as ancient in striata as the two types of MSPNs (Grillner & Robertson, 2016), their role in the basal ganglia circuit is less well conserved (e.g., Guirado et al., 1999), which, along with their rarity (always less than 5% of striatal neurons), suggests that they play a less vital role in representational functions of striata. In fact, the very small numbers of ChINs compared to D2-ENK-MSPNs suggests that they cannot effectively mediate another robust feature of extinction never considered by Crossley et al.; this feature being that extinction is more context-delimited than the initial acquisition (Bouton et al., 2012; Delamater & Westbrook, 2014; Todd et al., 2014). This robust feature conflicts with use of synapses onto ChINs to mediate extinction learning. Because there are few ChINs, context coding via ChINs would be coarse, and discrimination poor, despite the extensive branching of ChINs being hypothetically sufficient to cover the whole of striatum. Equally problematic: because each ChIN projects to many MSPNs, learning due to extinction of one response in a context would generalize too broadly, by depressing many other responses in the same context. Neither problem attends our model. Because about 46% of striatal neurons are D2-MSPNs, extinction learning via synapses onto small subsets of them can be highly context-, cue-, and response-specific.

Unlike prior alternatives, the present model suggests that fast adaptive switching of bases for decision making can be achieved in a stochastic model by cumulative DA-dependent learning at thalamo- and cortico-striatal synapses. The centrality of DA-related asymmetries across striatal sectors to the model’s function is consistent with the hypothesis (Wickens et al., 2007) that DA clearance times at the synapse, which depend on DAT levels, may have important consequences for decision making. The independence of the mechanism presented here from cortical intervention, as well as its generalizable structure, provide a basis for large scale studies of how basal-ganglia function can co-evolve with pallium or neocortex in lineages whose life styles make use of different types of representations. The mechanisms simulated here can provide an arbitration mechanism for competing representations of unknown type and level of abstraction. The mechanism’s independence from specific input representations helps explain how it has been successful at mediating control of a huge range of behaviors including: locomotion, grasping by jaw or hand, attention and gaze, categorical speech perception, speech production, storage in working memory, etc.

A computational model presented in Keramati & Gutkin (2013) also used a VMS-DLS cascading learning rule, inspired by the VMS-DLS “spiral” anatomy reviewed in Haber et al. (2000). However, the model of Keramati & Gutkin (2013), unlike the current model, assumed that the asymptotic values of association strengths are *normally equal* along the VMS-DLS axis. They proposed that typical addictive drugs, such as cocaine, are habit forming because of their ability to engender an *abnormal* gradient of increasing weight asymptotes along the VMS-DLS axis (e.g., as reported by Willuhn et al., 2012). Such a model offers no explanation for the *normal* transitions into habit mode that are routinely seen in behavioral studies using non-addictive rewards. In contrast, our model assumes that there is an inherent gradient of asymptotic synaptic weight potentials along the VMS-DLS axis due to regional differences in DAT, dendritic branching, synaptic spine density, etc. (see Table 1). It can explain switches to habit mode in the absence of any drug-induced abnormality. Also, unlike the serial learning model in Keramati & Gutkin (2013), the current, parallel-learning, model does not require a delayed onset of DA release in DLS relative to VMS/DMS.

A key reason for treating the gradient of weight asymptotes as a normal feature is its support of an emergent SAT governed by the reward-earning values of cue-guided actions. As long as reward is reliably obtained, the system will transfer control to regions with higher weight saturation points, which correspond to regions with inputs based on faster-computed representations. This, combined with larger weights, guarantee faster behavior onset times. However, the process of transfer is self-regulating, because in order to properly interpret stimulus and mnemonic information pertinent to the currently active action-outcome contingency, and choose a response actually able to obtain reward at high probability, some above-minimum processing complexity is often required. The push-pull relationship between speed and accuracy inherent to the transition mechanism of the present model provides a natural tendency for the system to converge *toward* an optimized stochastic mix of reaction times. However, the emergent speed-accuracy tradeoff need not correspond to the optimal strategy from a mathematical standpoint. Given the dependence of convergence on diverse factors (Table 1), e.g., genetically variable DA-signaling parameters, solutions are predicted to vary across individuals.

A simulated effect of cocaine was explored in Fig. 9, but several other pharmacological agents known to affect DAT and synaptic spine formation have yet to be assessed in the context of the model. Prominent among such candidates for modeling studies are amphetamines (Giros et al. 1996; Robinson & Kolb, 1997; Jedynak et al., 2007) and ethanol (Zhou et al. 2007; Rice et al. 2012; Ibias et al. 2015; Lovinger & Alvarez 2017). Other VMS to DLS gradients that deserve modeling study include the distributions of cannabinoid receptors (Herkenham 1991; Mailleux & Vanderhagen 1992; Davis et al., 2018)) and opioid receptors (Voorn et al. 1996; Daunais et al., 2001; Oude Ophuis et al. 2014). Although drugs and their withdrawal have diverse effects that go well beyond what can be represented in the current model, it offers a way to understand how their net effects on habit formation often depend on how they alter the rate and asymptote of learning in more than one striatal compartment. Because that includes core and shell regions of VMS, both striatal sectors offer promising targets for near-term elaborations of the model. Beyond drug effects, such an elaborated model could help bring neural data to bear on the evaluation of hypotheses such as the Trask et al. (2017) proposal that extinction reflects various underlying mechanisms, e.g., “occasion setting” of Pavlovian responses (mediated partly by VMS) versus direct inhibition of instrumental responses (mediated by DMS and DLS). Another priority is the phenomenon of Pavlovian-instrumental transfer (PIT), the robust finding that previously learned cue-reward or cue-pain associations can be leveraged, respectively, to boost or depress instrumental responses (review in Corbit & Balleine, 2016).

Although simulations showed that the current model can explain emergent speed-accuracy tradeoff (SAT) phenomena, recent studies (de Froment et al., 2014; Thura & Cisek, 2014) confirm the intuition that additional factors, such as persistent context stimuli, and the effort or pain associated with correct response, affect such tradeoffs. While effects of cue processing delays can be treated by the current model, by design it does not address many issues involving the persistent contexts, transient cues, and compounds or conjuncts thereof that may be represented in a given species. Also, although effort costs affect learning and decisions, and have some effects on DA release, much more is known about how benefits affect DA release than about how expected costs or pain do. Although more research will be needed to understand such effects, we predict that some of these effects will operate via forebrain gradients that affect two key DA-dependent variables: learning rate and asymptotic synaptic strength.

## Model and Methods

The model used to generate the preceding results consists of two structurally-similar compartments, operating in parallel, each of which represents a cortical-basal ganglia loop (Figure 3). Each compartment contains four distinct populations – a stimulus representation population, which activates in the presence of an external stimulus; an input/output population representing the cortical component of the cortical-basal ganglia loop, a GO pathway, and a NOGO pathway. The GO and NOGO pathways correspond to “direct” and “indirect” anatomical pathways through the basal ganglia; the direct pathway promotes responses, while the indirect pathway inhibits responses. Each pathway has cells of origin in the striatum, distinguished by their DA receptor expression. The direct (**GO**) pathway cells of origin are **D1-MSPNs** (Dopamine receptor type-1 medium spiny projection neurons), whereas the indirect (**NOGO**) pathway cells of origin are **D2-MSPNs** (Dopamine receptor type-2 medium spiny projection neurons.) The difference in function between DA receptors expressed in the these opposing pathway forms the empirical basis for learning rules that produce opponent weight adjustments while still using a shared learning signal (ie., burst and dips of dopamine release).

The representation of the cortical, thalamic, and basal ganglia circuits used for this simulation is a “lumped” representation: it omits several anatomical populations and connections that would normally be necessary relays through the system, and that have been included in some prior models (e.g., Brown et al., 2004; Civier et al., 2013). All of the omitted elements would be needed to model some physiological results not modeled here. For example, signal processing in the (omitted) pallidal and thalamic stages implies that basal ganglia control of thalamic activation of cortical plan representations works through disinhibition of thalamo-cortical loops (Brown et al., 2004) rather than by direct excitation. Also, the two layers of cortical neurons in Brown et al. (2004), replaced here by a single layer, would be needed to model tasks in which choices precede actual behavioral performance commitments by significant intervals, as well as some SAT effects and physiological observations, such as those recently reported by Thura & Cisek (2017), as they noted. Relatedly, reinforcement-guided weight adjustment, restricted in the current model to cortico-striatal synapses, was distributed among cortico-cortico and cortico-striatal synapses in the model of Brown et al. (2004). In summary, the present model uses an action-selection model with minimally-complicated interactions, in order to accentuate the essential simplicity with which dopaminergic and striatal asymmetries (omitted from Brown et al., 2004) may facilitate control switches that are important for habit formation and SAT. The simplifications have the advantage of allowing much clearer attributions of the effects we do model to particular mechanisms/parameters.

Each neural population used in this simulation is organized as a matrix with two dimensions: first, along the set of sensory stimuli available to the population; and second, along the set of possible behaviors. There are a total of four unique sensory stimuli available and three unique behavioral outputs. From these two sets, a population is generated with one cell which accounts for each combination of stimulus and action: Stimulus 1/Action 1; Stimulus 1/Action 2; and so on. This segregates each possible stimulus / action pair into its own channel through the cortical-basal ganglia loop, allowing individual synaptic weights to affect the potency of one and only one action / outcome pairing.

The model makes use of two distinct compartments, the “primary” and “secondary” compartment. The secondary compartment is identical in structure to the primary compartment, with one important difference: it has two extra rows of cells along the sensory-stimuli axis that can be stimulated. These extra rows are activated by stimuli 3 and 4, which by experimental design are only co-presented along with one of the two primary stimuli (stimulus 1 or 2). These so-called conjugate cells are intended to represent a richer variety of cortical processing available to cortices that innervate the ventromedially-located regions of the basal ganglia when compared to those that innervate the dorsolateral zone.

With the span of stimuli and actions for a given population, there are three layers that share the same number of elements and interact with one another within the population: the cortical (CTX), D1-type-striatal (GO), and D2-type-striatal (NOGO) layers. The CTX layer receives excitation as a direct result of sensory input and, when sufficiently activated, produces a motor output corresponding to the action represented by the maximally-activated CTX node. The GO layer provides direct positive feedback to the CTX population through a pair of excitatory weights, one of which (the CTX to GO weight) can be adjusted during learning; similarly, the NOGO layer provides net inhibitory feedback through one excitatory adjustable CTX-to-NOGO weight and one fixed inhibitory NOGO-to-CTX weight. Equations 1-3 (below) govern cell activations in these three layers.

To ensure a degree of randomness in the model’s decision-making, as required for exploratory learning, a stochastic component was incorporated at the cortical stage. This stochastic element consists of the addition of a noise input that was newly generated and applied at each time step to all CTX cells. These noise signals, randomly drawn from a Gaussian distribution, sufficed to cause noticeable variability in the responses generated, especially prior to conditioning, but also, to a reduced degree, after the adjustments to CTX-GO/NOGO weights that resulted from conditioning trials.

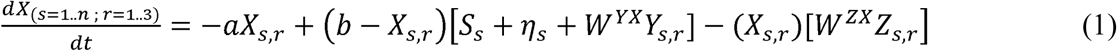

Equation 1 is the cortical (CTX) activation equation. Constants, passive decay term a=0.01, upper bound term b=1, and number of unique stimulus representations n = 2 for the primary compartment and n = 4 for the secondary compartment. Indices s = 1..n refer to stimulus 1, stimulus 2, etc. while indices r = 1..3 correspond to response 1, response 2, and response 3. S_s_ represents the input activation caused by the external stimulus and takes a value of either 0 or 1 for each s=1..n. *η*_*s*_ is a Gaussian random variable with mean 0 and standard deviation 0.1, generated de novo at each timestep for each s. Y_sr_ and Z_sr_ represent the activations of the GO and NOGO populations respectively, weighted by the identical weight terms W^YX^ and W^ZX^ ie., the weights from Y to X (GO to CTX) and Z to X (NOGO to CTX) respectively. These weights from the GO and NOGO pathway onto the CTX are fixed at a value of 0.5 in all simulations.

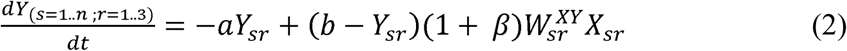

Equation 2 is the striatal GO neuron activation equation. Constants a=0.01, b=1, n = 2 (primary compartment) or n = 4 (secondary compartment). These equations share the basic form of Equation 1, with identical constants. The weight terms 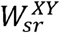 represent the adaptive weights from Xsr to Ysr (CTX to GO), which may take on different weight values between 0 and 1 for each individual s,r pair. These weights are each initialized with a value of 0.25. Parameter β had a value of 0 in both compartments for the most simulations; however, in the simulations producing Figures 14 and 15, β had a value of 0.2 in the secondary compartment, and β had a value of 0.1 in the primary compartment only during the simulations summarized in Figure 15.

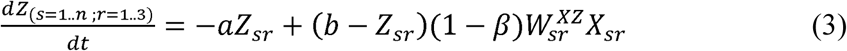

**Figure 15.**
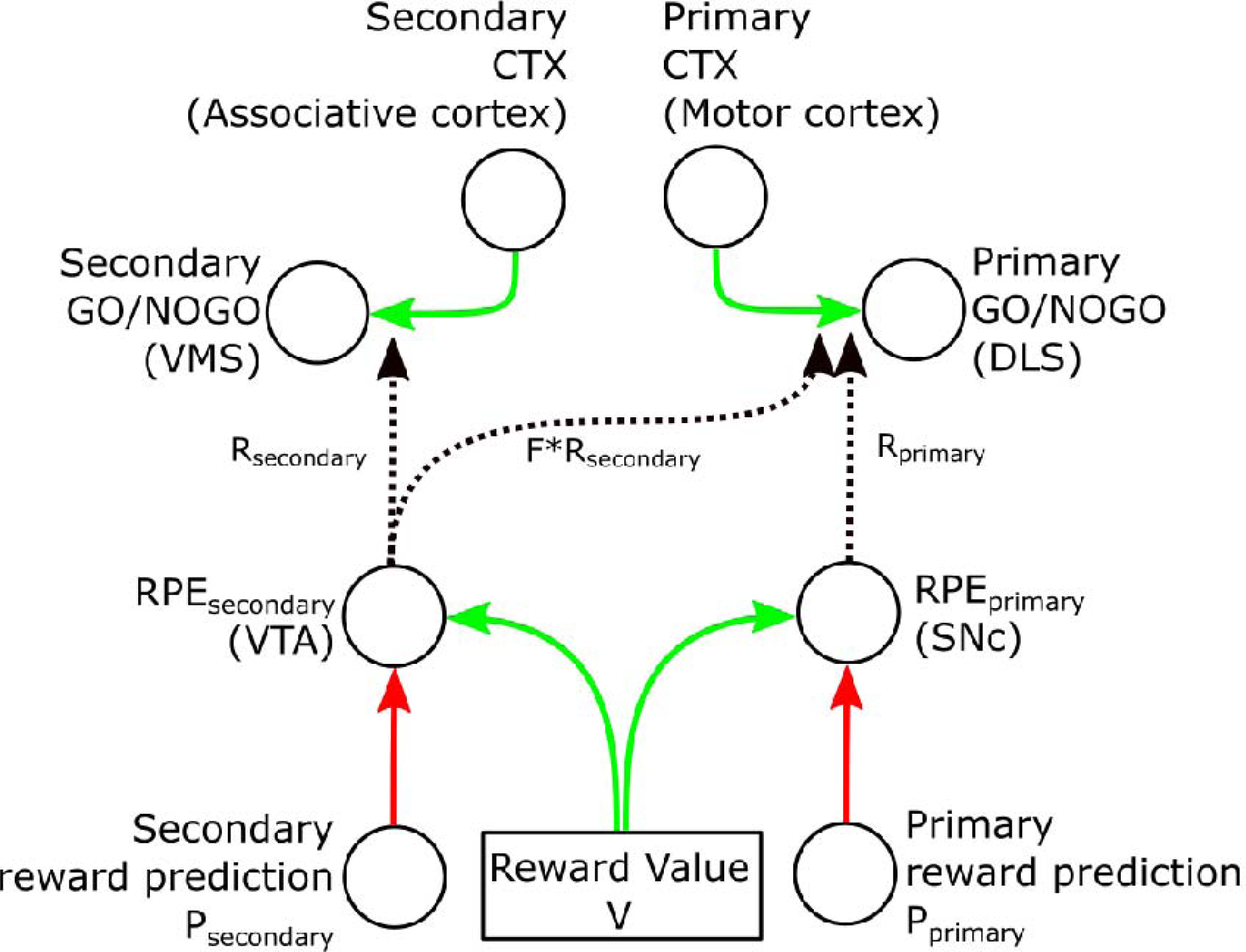
Reinforcement system diagram. This figure illustrates the set of factors that contribute to the final value of the reinforcement learning signal that is used to guide synaptic weight changes in the model. The prediction signal, P, which differs between compartments (P_primary_ and P_secondary_) is used to determine RPE values, which are then used to compute signal R. The reward value, V, depends on the behavioral output of the model, the contingency, and the pathway (GO or NOGO), per Table 5. These signals combine to adjust the weights between the CTX and GO/NOGO layers according to the rule given by Equation 7.

**Figure 16.**
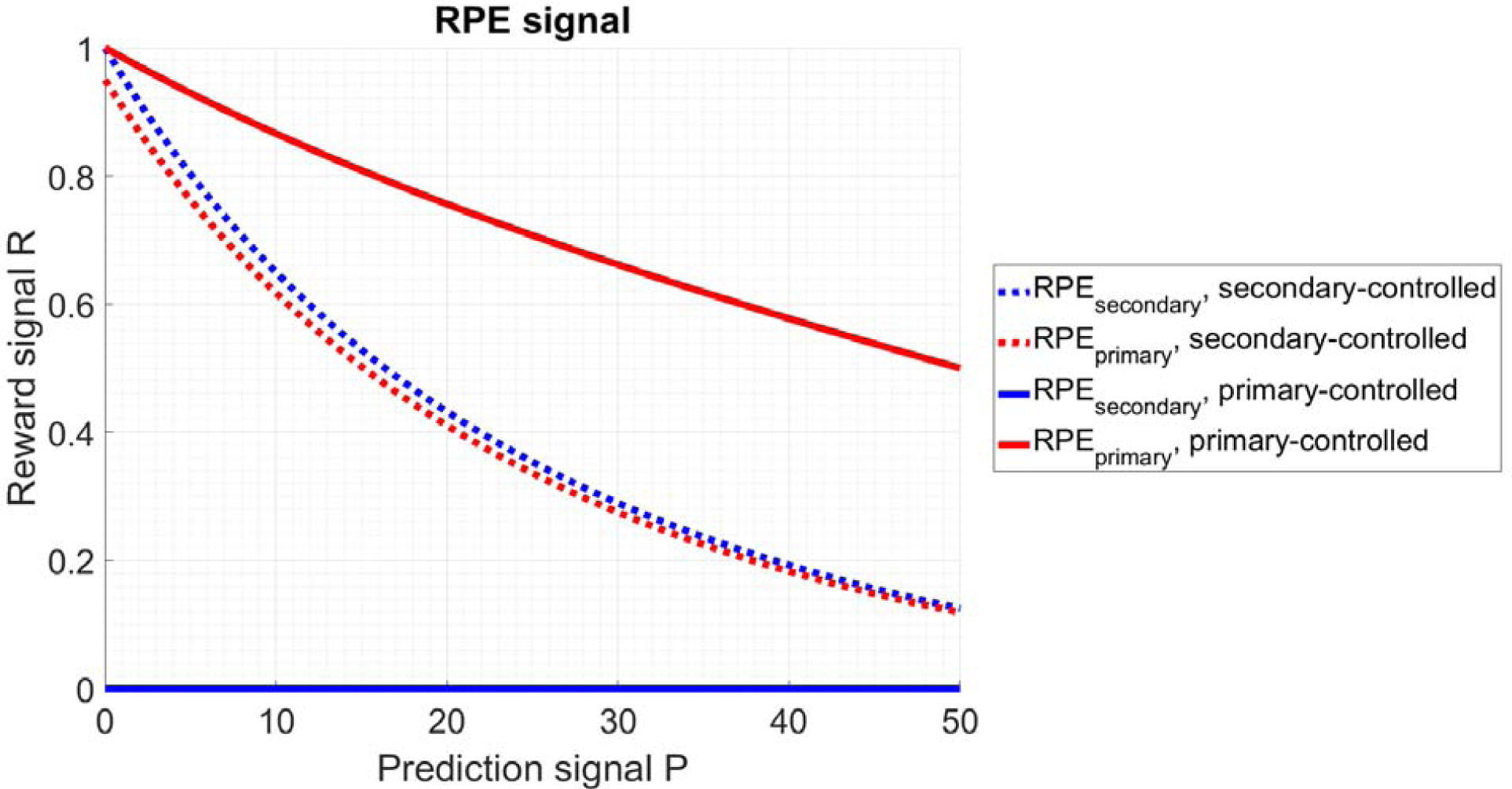
Reward prediction error signals RPE_secondary_ and RPE_primary_ (Eqns. 5 and 6) as a function of reward prediction P (Eqn. 4). The above graph depicts the RPE signals obtained following a trial ending in reward. In the case of a reward obtained following a secondary controlled response, both compartments receive an RPE signal determined by the secondary prediction signal RPE_secondary_ (dotted blue line); the primary RPE (dotted red line) is a copy of the secondary RPE, scaled by a factor of F (F=0.95 in this graph for visual clarity; however in all simulations shown in the Results section, F=1, meaning both RPE_primary_ and RPE_secondary_ take the same value). In the case of a reward following a primary controlled trial, the secondary compartment receives no RPE signal (solid blue line) and therefore no weight adjustment, while the primary compartment receives an RPE signal (solid red line) that is larger at any value of P than the comparable RPE signal used in the case of a secondary-controlled trial, reflecting the reduced prediction effects observed in experimental DLS dopamine release events following the acquisition of a predictable reward. After 50 identical reward outcomes, predictability index P reaches a maximum value of 50, and both RPE curves reach their minimum value. Thus the model’s approximation for computing RPEs only accounts for the past 50 trials.

Equation 3 is the striatal NOGO neuron activation equation. Constants a=0.01, b=1, n = 2 (primary compartment) or n = 4 (secondary compartment). These equations share the basic form of Equation 1, with identical constants. The GO and NOGO pathways have distinct weight values 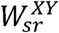 and 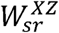 respectively, and are adjusted according to opposite learning rules; 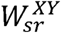, the GO or D1-MSPN synaptic weight, is always adjusted opposite to 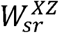, the NOGO or D2-type synaptic weight. The parameter β had a value of 0 in both compartments for most simulations; however, in the simulations producing Figures 13 and 14, β had a value of 0.2 in the secondary compartment, and β had a value of 0.1 in the primary compartment only during the simulations summarized in Figure 14.

At the beginning of each trial, activities in all neural populations are reset to initial conditions, though the matrix of weights retains any alteration that occurred due to learning in prior trials. Neural activities are set to a baseline of 25% maximum activation (0.25). A stimulus is then provided to the CTX layer of both compartments according to the currently active experimental paradigm, for up to 300 time steps, at which point the trial ends with no action performed unless one of the CTX neurons in either compartment has become sufficiently active to elicit behavior. Behaviors are produced when a neuron in the CTX layer exceeds a globally defined activation threshold, with this value being set to 75% maximum activation (0.75).

If any CTX neuron exceeds the threshold to produce a behavior at any time step, the trial is immediately ended, interrupting any further accumulation of activation. Once this threshold crossing occurs, the behavior corresponding to the most-active CTX neuron (according to the scheme shown in Tables 1 and 2) is produced and a reward value is determined based on the currently-active contingency. In the event of a tie, a winner is chosen at random from among the tied options. The random tiebreaker is typically only used once per learning regimen to destabilize neurons activated in lockstep due to (rare) identical weight initializations. Initial weights are set to 20% of maximum (0.2) plus or minus an initial offset between the GO and NOGO weights (0.1 in secondary compartment, 0.02 in primary compartment) so that the initial weights for the secondary compartment are 0.1 (NOGO) and 0.3 (GO) in the secondary compartment; and 0.18 (NOGO) and 0.22 (GO) in the primary compartment. These initial weights are perturbed by a small amount of noise (±0.02) in order to reduce the number of “tiebreaker” trials and produce a more uniform initial acquisition curve. Table 6 presents an example matrix of initial GO weights, and Table 7 presents an example of how that matrix might have changed due to learning trials with a contingency that rewarded response 1 in the context of stimulus one.

**Table 3.** Organization of primary compartment populations.

|  |  |  |
| --- | --- | --- |
| Stimulus 1 / Action 1 | Stimulus 1 / Action 2 | Stimulus 1/ Action 3 |
| Stimulus 2 / Action 1 | Stimulus 2 / Action 2 | Stimulus 2/ Action 3 |

**Table 4.** Organization of secondary compartment populations.

|  |  |  |
| --- | --- | --- |
| Stimulus 1 / Action 1 | Stimulus 1 / Action 2 | Stimulus 1/ Action 3 |
| Stimulus 2 / Action 1 | Stimulus 2 / Action 2 | Stimulus 2/ Action 3 |
| Stimulus 3/ Action 1 | Stimulus 3 / Action 2 | Stimulus 3/ Action 3 |
| Stimulus 4/ Action 1 | Stimulus 4 / Action 2 | Stimulus 4 / Action 3 |

**Table 5.** Experimental paradigms used in results section. This table shows the stimulus-response contingencies pertaining to each Results figure, along with the number of trials presented of each. This table is intended as a companion to the text accompanying each figure, which provides added detail about each paradigm.

| **Figure number** | **Stimulus / Response contingency leading to reward; trials active (stimuli presented)** |
| --- | --- |
| 5 (5a, 5b, and 5d) | Stimulus 1 / Response 1; 100 (Stimulus 1)<br>No stimulus / response (extinction); 100 (Stimulus 1)<br>Stimulus 1 / Response 1; 100 (Stimulus 1) |
| 6 | Stimulus 1 / Response 1; 500 (Stimulus 1)<br>Stimulus 1 / Response 2; 500 (Stimulus 1)<br>Stimulus 1 / Response 1; 500 (Stimulus 1)<br>Stimulus 1 / Response 2; 500 (Stimulus 1)<br>Stimulus 1 / Response 1; 500 (Stimulus 1)<br>Stimulus 1 / Response 2; 500 (Stimulus 1) |
| 7 | Stimulus 3 / Response 1, Stimulus 4 / Response 2; 50 (50% Stimulus 3, 50% Stimulus 4)<br>Stimulus 3 + Stimulus 1 / Response 1; 100 (Stimuli 1 and 3)<br>Stimulus 4 + Stimulus 1 / Response 2; 50 (Stimuli 1 and 4)<br>Stimulus 3 / Response 1, Stimulus 4 / Response 2; 50 (50% Stimulus 3, 50% Stimulus 4)<br>Stimulus 3 + Stimulus 1 / Response 1; 1000 (Stimuli 1 and 3)<br>Stimulus 4 + Stimulus 1 / Response 2; 50 (Stimuli 1 and 4)<br>Stimulus 3 / Response 1, Stimulus 4 / Response 2; 50 (50% Stimulus 3, 50% Stimulus 4)<br>Stimulus 3 + Stimulus 1 / Response 1; 2000 (Stimulus 1 and 3)<br>Stimulus 4 + Stimulus 1 / Response 2; 50 (Stimulus 1 and 4) |
| 8 | Stimulus 1 or 2 / Response 1; 1500 (50% Stimulus 1, 50% Stimulus 2)<br>Stimulus 3 or 4 / Response 2; 1500 (50% Stimulus 3, 50% Stimulus 4) |
| 9 | Stimulus 1 / Response 1; 500 (Stimulus 1)<br>Stimulus 1 / Response 2; 500 (Stimulus 1)<br>Stimulus 1 / Response 1; 500 (Stimulus 1)<br>Stimulus 1 / Response 2; 500 (Stimulus 1)<br>Stimulus 1 / Response 1; 500 (Stimulus 1) |
| 10 | Stimulus 1 and 3 / Response 3, Stimulus 2 / Response 2 (10% chance); 2000 (70% Stimulus 1, 30% Stimulus 2, 50% Stimulus 3) |
| 12, 13, 14 | Stimulus 1 / Response 1; 500 (Stimulus 1)<br>Stimulus 1 / Response 2; 500 (Stimulus 1)<br>Stimulus 1 / Response 1; 500 (Stimulus 1)<br>Stimulus 1 / Response 2; 500 (Stimulus 1)<br>Stimulus 1 / Response 1; 500 (Stimulus 1)<br>Stimulus 1 / Response 2; 500 (Stimulus 1) |

**Table 6.** Example primary compartment CTX-to-GO weight matrix 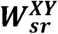 initial conditions, before training. All weight values are within the bound [0.45 0.55].

|  |  |  |  |
| --- | --- | --- | --- |
| *Stimulus 1* | **0.47** | **0.52** | **0.55** |
| *Stimulus 2* | **0.54** | **0.5** | **0.42** |

**Table 7.** Example primary compartment CTX-to-GO weight matrix 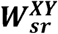 values after training. Weights shown are for demonstration purposes and were not generated through simulation. The weight values shown illustrate the adjustments made during a training period in which the contingency Stimulus 1 → Response 1 is rewarded and other outcomes are devalued. In this scenario, assuming the NOGO pathway remains fixed, weights will be decremented in order of value until the weight corresponding to the rewarded contingency is highest, at which point reward can be consistently achieved until the weight limit is reached (for the primary compartment, Γ=0.9). Weights below the initial value of the rewarded contingency (e.g., S2/R3 above = 0.42) are not adjusted.

|  |  |  |  |
| --- | --- | --- | --- |
| *Stimulus 1* | **0.9** | **0.46** | **0.46** |
| *Stimulus 2* | **0.46** | **0.46** | **0.42** |

Weights in the model are adjusted according to a piecewise linear learning rule, specified below in Eqn. 7. However, across two different compartments (each of which is competing for control of behavior using a separate set of CTX, GO, and NOGO cells) two parameters of this equation differ: the learning rate λ and threshold limit Γ. Variation between these terms leads to crossing points in learned weight trajectories between populations, causing the control of behavior to be exchanged without destructively overwriting existing weights.

The difference in how reward affects the GO and NOGO synapses in the secondary versus primary compartments creates a functional transition point for behavior in some cases of extended training. Given sufficient rewarded trials, control of behavior transitions from the secondary compartment-having a high learning rate λ but low saturation point Γ - to the primary compartment, which continues to increment its weights (and thus reduce the amount of time needed to exceed activation threshold) during late learning trials as a result of its high saturation point Γ. Figure 2 depicts the weight cross-over that mediates such a transition.

Assuming the primary compartment is able to reliably achieve a rewarded outcome, it will maintain control of behavior after the behavioral switching point is crossed. If, however, the information available to the primary compartment is often insufficient for it to choose a response that earns reward, then control of behavior will quickly transition back to the secondary compartment. In such cases, the weights in the two compartments can maintain an equilibrium across multiple trials and remain at or near the crossing point.

At the end of each trial, reinforcement is applied to the synaptic weights connecting CTX to GO cells as well as the weights connecting CTX to NOGO cells. Reinforcement is applied only once per trial, and the effect at the synapse can be either facilitative or depressive. Weight adjustments in the GO and NOGO pathways are always opponent, with a facilitative effect on the CTX-GO synapse being paired to a depressive effect on the CTX-NOGO synapse. Rewarded responses produce reinforcement that strengthens the GO pathway while weakening the NOGO pathway, with the accompanying change in the balance of excitation and inhibition implying their CTX target will accumulate activation more rapidly on the next trial following positive reinforcement. Likewise, unrewarded responses weaken the GO pathway but strengthen the NOGO pathway, causing CTX to accumulate activation more slowly on following trials. While these changes are always opposite in sign, the magnitudes are not equal, with the NOGO pathway having a larger change in weight (150%) relative to the GO pathway. Negative reinforcement – which occurs if an incorrect decision is made – has twice the impact on weight adjustment as positive reinforcement. Anatomical and physiological data corroborate such asymmetries between aversive and appetitive conditioning in the two pathways (Lammel et al. (2011), Hikida et al. (2010)]. This relationship between the valence of reinforcement and its relative effects on the GO and NOGO pathway is depicted in Table 8.

**Table 8.** Reinforcement as a function of reward and pathway. In the biological reinforcement system, dopamine-releasing cells burst following the acquisition of reward and pause following omission of reward. The action of dopamine at the cortical-striatal synapse differs based on the receptor type expressed by their striatal targets: D1-type cells strengthen their synapse in response to a dopamine burst, while D2-type cells strengthen the synapse in response to a pause. This 2-factor relationship between reinforcement and synaptic weight change is captured in the table above: the GO pathway strengthens its synapses following a reward, while the NOGO pathway weakens them, with the opposite being true following an unrewarded trial. The relationships between these parameters are not equal in magnitude: the NOGO pathway is adjusted by 400% more than the GO pathway in both the rewarded and unrewarded case. Differences in the relative effectiveness of reinforcement enables learning-related changes in behavior to persist over multiple acquisition/extinction sequences.

| Value of Reinforcement, V | GO pathway | NOGO pathway |
| --- | --- | --- |
| Rewarded trial | V = 1 | V = -5 |
| Unrewarded trial | V = -1 | V = 5 |

Schultz et al. (1998) summarized data showing that as learning proceeds, dopamine neurons adjust their reward-related firing in such a way that their bursts are well characterized as a signal that scales with reward prediction error (RPE). If cues and responses reliably predict reward, this can be learned and the result is that the RPE signal – the reward-related release of dopamine -- is greatly diminished as trials accumulate. This trend is coarsely approximated by computations specified by Equations 4-6. First, equation 4 computes a sum, P, of 50 numbers as an index of the predictability of positive or negative outcomes of the latest 50 trials. The current reward outcome, V(t), is compared to with prior outcomes via a function *ε*(*V*(*t*), *V*(*t*–∂)), which evaluates as 1 if the current reward is identical to the reward obtained δ trials ago, and 0 otherwise. For example, a sum of 50 results if the last 50 outcomes are all identical to the current one, but a sum of 25 results if V alternated between positive reward (V=1) and negative reward (V=-1) across those 50 trials. Higher values of P cause greater reductions in RPE signals, according to Equations 5 and 6.

The calculation of the reward prediction error signals differs depending on the associated compartment. Based on the observations that ventral striatum responses to reward and RPE adjustments are more dramatic than those in DLS (Apicela et al. 1991; Schultz et al. 1992; Parker et al. 2016), the RPE curve applied to the secondary compartment, RPE_secondary_ has a steeper dropoff than that of the primary compartment, RPE_pri__mary_. Both equations 5 and 6 capture a power-law relationship between the prediction signal P and the ratio of unexpected rewards to expected rewards, but with powers chosen to ensure that the model’s RPE signal for the secondary compartment has a steeper dropoff with P than that for the primary compartment. Beyond making the model respect biologically observed differences in RPE behavior, this use of separate RPE curves for the secondary and primary compartments enhances the ability of over-training to cause transitions of control by the primary compartment, relative to what would be obtained with equivalent RPE signals. As with equation 4, these equations are intended as coarse approximations of biological reward prediction phenomena.

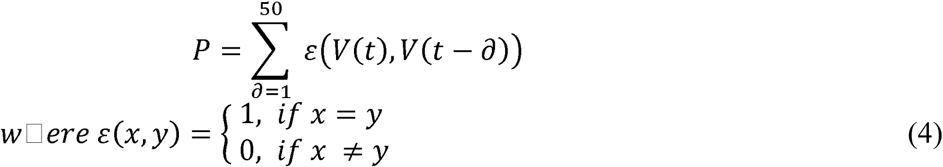

**Equation 4.** Outcome predictability, P. The value of P is determined by the history of reward value V obtained according to Table 3. For each of the prior 50 trials, the current trial’s reward value is checked against the reward value obtained in each of the previous trials. P is the sum total of matching reward values across the last 50 trials.

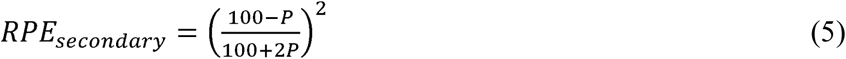

**Equation 5.** Secondary compartment reward prediction error signal RPE_secondary_. The reward prediction error signal is calculated parametrically from the outcome predictability value P. This reduces the RPE, and thus the impact of a reward on learning, more with each matching reward value, to a minimum of about 5% of its nominal value at 50 matches.

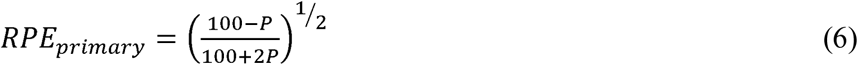

**Equation 6.** Primary compartment reward prediction error signal RPE_primary_. The reward prediction error signal is calculated parametrically from the reward prediction value, P. This reduces the RPE, and thus the impact of a reward on learning, more with each matching reward value, to a minimum of about 50% of its nominal value at 50 matches.

After each trial, a reinforcement signal occurs in both the primary and secondary compartments. Depending on which compartment gained control of behavior, however, the relative magnitudes of these reinforcements differ, in such a way that the compartment that was in control receives a larger reinforcement value than the inactive compartment. The basis for this spreading of reinforcement between the two compartments is the anatomical observation of collaterals from dopaminergic neurons to the cortical-striatal synapses of nearby sub-regions. Haber (2000) noted that these projections preferentially project in a “spiral” along the VM-DL axis, leading to our choice of parameterization for a much stronger secondary-to-primary spreading of reinforcement than vice versa. A feed-forward scaling factor, F = 1, is applied to rewards obtained under secondary compartment control when applied to reward impact on the primary compartment; however, there is no reciprocal application of reward obtained under primary control to the secondary compartment (see Table 4; Figure 3). Based on the observation that dopamine release events in VMS may be of comparatively lower magnitude than those in DLS (Cragg 2004; Calipari 2012), values of F lower than 1 are biologically plausible; lower values of F correspond to an extended number of secondary controlled trials before the transition to primary control occurs.

The effects of reinforcement on the synaptic weights of CTX with both the GO and NOGO pathways differ between the primary and secondary compartment. These differences arise from two parameters, λ and γ, which correspond, respectively, to the learning rate and upper weight limit represented graphically by Figure 3. Thus the primary compartment has a relatively higher weight limit γ when compared to the secondary compartment γ, but the primary compartment’s learning rate λ is lower than the λ of the secondary compartment. Both compartments have a lower weight limit of 0. The piecewise linear learning rule, and its incorporation of these two parameters, is given in Equation 7.

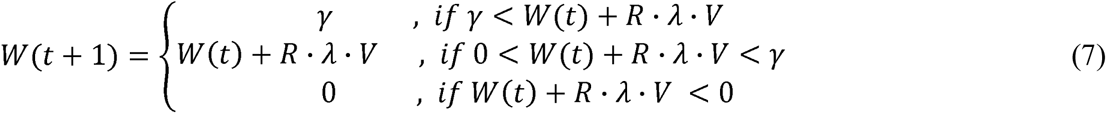

**Equation 7. Learning equation.** This piecewise linear learning equation produces adjustments to synaptic weights in both the secondary and primary compartments, based on the reinforcement values R_secondary_ and R_primary_, obtained according to Table 9, after each trial ends. The parameter λ represents the overall rate of learning, while parameter γ sets the upper asymptotic limit of weight value. λ and γ are parameters that vary inversely between the two compartments - the deliberative compartment has high λ and low γ, and vice versa for the habitual compartment. Within the primary compartment, λ = 0.0036 and γ = 1; in the secondary compartment, λ = 0.0045 and γ = 0.35. R is a variable that takes on the value of R_primary_ in the primary compartment and R_secondary_ in the secondary compartment. GO pathway weights are adjusted oppositely to those in the NOGO pathway, according to the value of ***V***, which takes values 1 or −1 in GO pathways and 5 or −5 in NOGO pathways, depending on the outcome (reward or not), as specified in Table 6.

**Table 9.** Asymmetrical projection of reinforcement signals to the two compartments. Although only one compartment can determine behavior on each trial, a reward obtained as a result of the correct response can lead to a reinforcement signal that is distributed to both compartments. Based on the values of their respective RPE signals, values of R for the secondary and primary compartment are obtained. This final scaled version of R is the value used for weight adjustment according to Equation 7. Reinforcement signals consequent to behavior controlled by the secondary compartment reached both it and the primary compartment (F = 1), but not the reverse. This reflects neuroanatomical data showing that dopaminergic axons collateralize in the VM to DL direction but not the reverse (Haber, 2000), as well the functional hypothesis that the input signals needed to excite DA neurons that project to the secondary compartment (DMS) are absent when signals that might otherwise activate its loop are curtailed as soon as a (by its nature, *faster*) choice is made by the primary compartment.

| Asymmetrical projection to determine total reinforcement, $R$ | Reinforcement in secondary and primary compartments |
| --- | --- |
| Secondary-controlled trial | $R_{\text{secondary}} = RPE_{\text{secondary}}$<br>$R_{\text{primary}} = F * RPE_{\text{secondary}}$ |
| Primary-controlled trial | $R_{\text{secondary}} = 0$<br>$R_{\text{primary}} = RPE_{\text{primary}}$ |

Because the primary compartment has access to less complete stimulus information, it is unlikely that it would be sufficient to produce a rewarded action if the secondary compartment had already failed to do so with its relatively superior information about the environment. Coupling its learning to that of the secondary compartment allows the primary compartment to produce the same action that the secondary compartment has been previously responsible for producing, but *with a reduction in production time* due to the higher cortical-basal ganglia weight threshold and simpler stimulus input in the secondary compartment. As long as a stimulus in the environment that is accessible to the primary compartment remains sufficiently predictive of reward, the primary compartment will retain control with a reduced production time; however, if the complex stimulus processing accessible to the secondary compartment is necessary to predict the rewarded outcome, repeated failures by the primary compartment to produce reward will cause control to revert to the secondary compartment.

